# Age-related cognitive decline in house crickets reveals conserved patterns of sensory and learning deficits across the lifespan

**DOI:** 10.1101/2025.08.14.670304

**Authors:** Gerald Yu Liao, Jenna Klug, Warren Ladiges

**Author notes:** **Corresponding author.** Warren Ladiges.

## Abstract

Cognitive decline with age is characterized by impairments in learning, sensory discrimination, and decision-making. While mammalian models have advanced understanding of the neural substrates of aging, their use in large-scale behavioral studies is limited.

Invertebrate models, such as the house cricket (*Acheta domesticus*), offer short lifespans, high throughput, and conserved neurobiological pathways but remain underexplored in geroscience.

We developed a dual behavioral paradigm integrating an olfactory discrimination Y-maze and an escape learning task requiring crickets to override innate odor preferences. Adult, mid-age, and geriatric crickets were tested for sensory discrimination, associative learning, and decision speed. Morphological traits, including antennal and femoral metrics, were quantified to evaluate their influence on cognitive outcomes. Data were analyzed using ANOVA, ANCOVA, and logistic regression models.

Aging impaired olfactory preference and learning success, with geriatric crickets showing reduced task acquisition and memory retention. Mid-age individuals exhibited the slowest decision-making, suggesting an early onset shift in behavioral strategy. Morphological traits predicted aspects of sensory performance and physiological resilience, such as reduced weight loss in crickets with larger femoral dimensions but did not explain age-related cognitive deficits. Olfactory decline was particularly pronounced in males, mirroring sex differences observed in human cognitive aging.

House crickets exhibit hallmark features of cognitive aging, including sensory decline, learning impairments, and reduced resilience, independent of morphological deterioration. These findings establish the house cricket as a scalable invertebrate model for dissecting conserved mechanisms of neural aging and testing interventions to promote cognitive health.

## Introduction

Cognitive decline is a defining feature of aging, characterized by progressive losses in learning capacity, sensory discrimination, and decision-making speed [1-3]. Mammalian models have been central to identifying the neural correlates of cognitive aging, revealing alterations in synaptic plasticity, neuromodulation, and circuit dynamics [4-5], yet present inherent limitations for lifespan studies and large-scale behavioral assays. Invertebrate systems offer short lifespans, conserved neurobiological mechanisms, and experimental scalability, but remain underutilized in aging research [6-7].

The house cricket (*Acheta domesticus*) offers a promising model for investigating the biology of aging. It combines ecological validity with experimental tractability, owing to a relatively short lifespan, consistent behavioral responses, and anatomically accessible neural structures [8-9]. Central to its cognitive function are the mushroom bodies, which are paired neuropil structures containing two types of neurons (intrinsic Kenyon cells and extrinsic neurons), that support learning, memory and sensory integration, and are structurally and functionally analogous to the mammalian hippocampus [10]. These regions support adult neurogenesis (AN) throughout the lifespan, a process sensitive to both sensory input and age-related decline [11-12]. This makes the cricket especially well-suited for dissecting how aging influences cognitive performance at both behavioral and cellular levels.

Olfactory processing is particularly tied to mushroom body plasticity and cholinergic signaling [13]. In the house cricket, olfactory receptor neurons project via the antennal nerve into the antennal lobes, which relay information to the calyces of the mushroom bodies through well-defined projection neurons [14]. This pathway provides a direct anatomical and functional link between olfactory discrimination and higher-order learning centers in the insect brain. In humans, impaired olfaction is an early and reliable predictor of neurodegenerative disease onset and correlates with structural brain changes in memory-related regions [15-17]. Similar patterns of age-associated olfactory decline have been reported in insects [18], yet its relationship to sensory discrimination and learning remains poorly understood.

An additional advantage of the cricket model lies in its externally visible and quantifiable morphological features. Traits such as antennal length, body size, and mass vary across age and may influence perception or movement, providing an opportunity to dissociate true cognitive impairments from age-related changes in sensory mechanics or motor coordination [19]. This ability to link morphology to function adds further dimensionality to the cricket as a model for aging studies.

To investigate how aging alters sensory processing and associative learning, we developed a two-part behavioral paradigm combining an olfactory discrimination assay with an escape-motivated task. The latter leverages the cricket’s innate aversion to well-lit environments and builds on prior work linking such escape behaviors to neurogenesis and memory function [20]. By integrating olfactory cues into this ecologically grounded task, we assessed how crickets of different ages learn to associate sensory input with escape outcomes and whether their ability to discriminate between odorants declines with age.

We hypothesize that aging in house crickets impairs both associative learning and olfactory discrimination, reflecting deficits in memory formation, attention, and behavioral flexibility. By establishing and validating these paradigms in the house cricket, we aim to uncover behavioral signatures of cognitive aging in an invertebrate model and lay the groundwork for future studies targeting mechanisms of neural resilience and cognitive preservation.

## Materials and Methods

### Animal rearing and housing

House crickets were sourced from a national supplier (Fluker Farms Inc, Louisiana, USA) and maintained under controlled laboratory conditions designed for physiological stability and ethologically relevant behaviors [19]. The colony was genetically heterogeneous, originating from multiple regional strains across the United States.

Environmental parameters were held at 29 ± 1°C and 32 ± 3% relative humidity [21-22]. Crickets were housed in custom dual-layer Plexiglass enclosures: the outer layer provided insulation and containment, while the inner modular compartment allowed manipulation of light exposure. A continuous light schedule (24-h light) was used, with diagonally stacked egg cartons and opaque paper along the lower walls to create shaded microenvironments. This configuration enabled self-regulated light exposure, promoting natural phototaxis and thermoregulatory behaviors. Enclosures housed 20–30 individuals (10–15 cm^2^ per cricket), balancing space for movement with opportunities for social interaction [23]. Males and females were co-housed to preserve natural social structure. Pathogen exclusion protocols were not applied, maintaining environmental microbiota that may influence to aging phenotypes [24].

### Diet preparation and experimental design

Crickets were fed a standardized laboratory diet (Picolab Rodent Diet 20, 5053, Irradiated; Purina Mills, USA). To ensure uniform nutrient intake and minimize batch variability, powdered chow was incorporated into a gelatin-based matrix: gelatin was dissolved in distilled water at 100°C, cooled to 60°C, mixed with chow to form a homogenous paste, refrigerated overnight (4°C), and dehydrated using a commercial food dehydrator to prevent microbial growth. The dehydrated product was homogenized into a fine, uniform texture using a food processor and provided *ad libitum*. Hydration was maintained through water gel packs (Napa Nectar Plus, Systems Engineering Lab Group, Napa, CA), replaced every 48 hours.

At 3 weeks of age, 10 males and 10 females per group were assigned to one of three chronological age cohorts: adult (6 weeks), mid-age (8 weeks), or geriatric (10 weeks). All crickets underwent at least one week of acclimation to laboratory housing before testing, eliminating transport- or cage-induced behavioral artifacts. Thus, juvenile-assigned individuals were tested after one week, while geriatric-assigned individuals were aged under standard conditions for seven weeks prior to testing. This design equalized environmental exposure across cohorts and allowed behavioral phenotypes to stabilize, consistent with prior evidence that locomotor and cognitive behaviors normalize following sufficient housing stability [25].

Animals were monitored daily for health and survival; deceased individuals were promptly removed to prevent contamination and cannibalism. Behavioral testing began when individuals reached their assigned age threshold, under the environmental conditions described in the housing protocol. Each cricket was weighed at baseline and after behavioral testing using a precision analytical balance (Mettler Toledo XPR205, Switzerland). Changes in body mass were analyzed as an auxiliary measure of physiological stress or early frailty [26-27].

### Y-maze design

Olfactory discrimination was assessed in a custom 3D-printed Y-maze with three arms (12 cm x 5 cm) oriented at 120° around a central decision zone [28-29]. Walls (9 cm high) prevented escape while permitting unobstructed exploration. The maze was designed in OnShape CAD software, exported as STL files, sliced in PrusaSlicer, and printed on a Prusa i3 MK3S+ using 1.75 mm black PLA filament (0.2 mm layer height, 20% infill, bed 60°C, nozzle 215°C). This process ensured reproducible geometry and optimal resolution (Figure 1).

**Figure 1.**
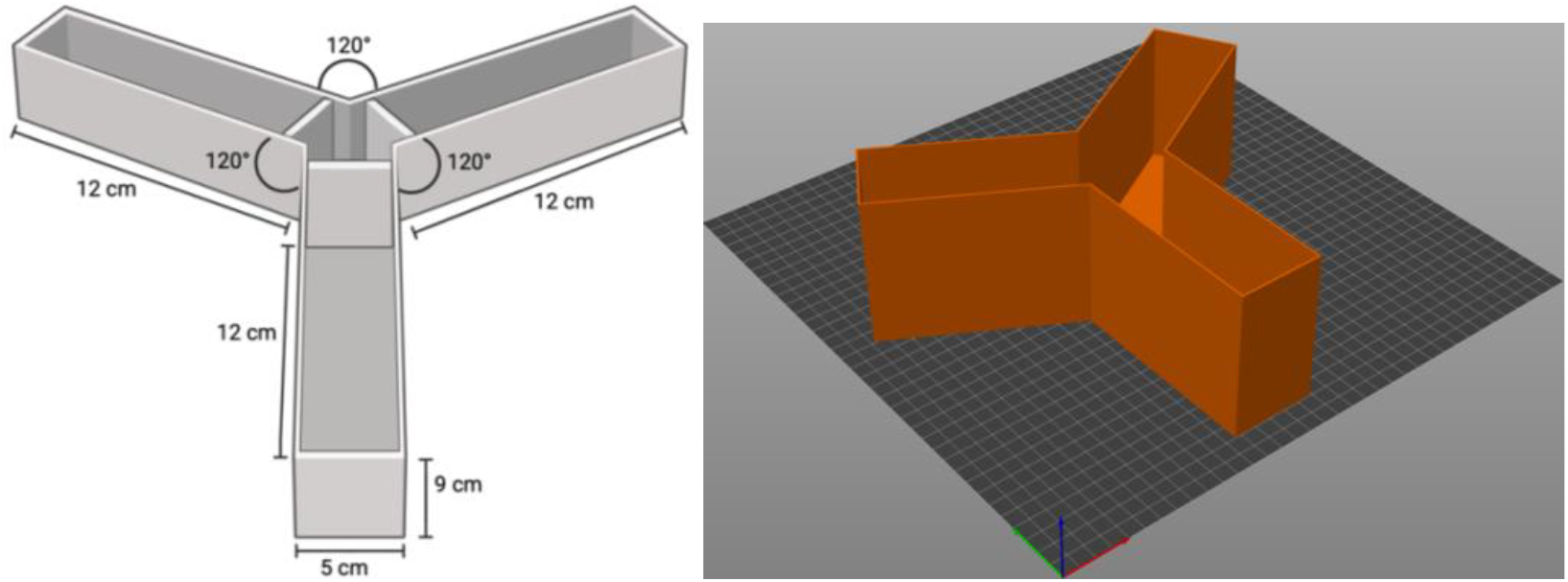
Schematic diagram of Y-maze constructed for cricket use, created using BioRender. com (left). Three-dimensional depiction of constructed Y-maze apparatus in OnShape (right).

To establish a controlled olfactory contrst, one choice arm contained vanilla odorant (1% vanilla extract, 1:1 diluted in distilled water), selected as an attractive stimulus based on empirical evidence of innate approach behavior in crickets [20]. The other arm contained cinnamon odorant (1% cinnamon extract, 1:1 diluted), chosen as a putative repellent based on entomology literature demonstrating cinnamaldehyde-induced repellency and behavioral disruption across insect taxa [30].

For each trial, a cricket was placed in the start arm and confined by a removable barrier for 10 s before release. Exploration was allowed for up to 30 s, with a choice recorded when the entire body entered a choice arm. The barrier was then replaced for 15 s to maintain odor exposure. Odorant positions were alternated between trials to control for spatial bias and assess path-chasing behavior, an indicator of odor tracking and retention (Figure 2). If no choice occurred within 30 s, the trial was excluded. Each cricket completed eight consecutive trials with 30 s inter-trial interval to minimize residual odor cues. The primary outcome was the percentage of vanilla arm entries as a measure of odor preference and decision-making.

**Figure 2.**
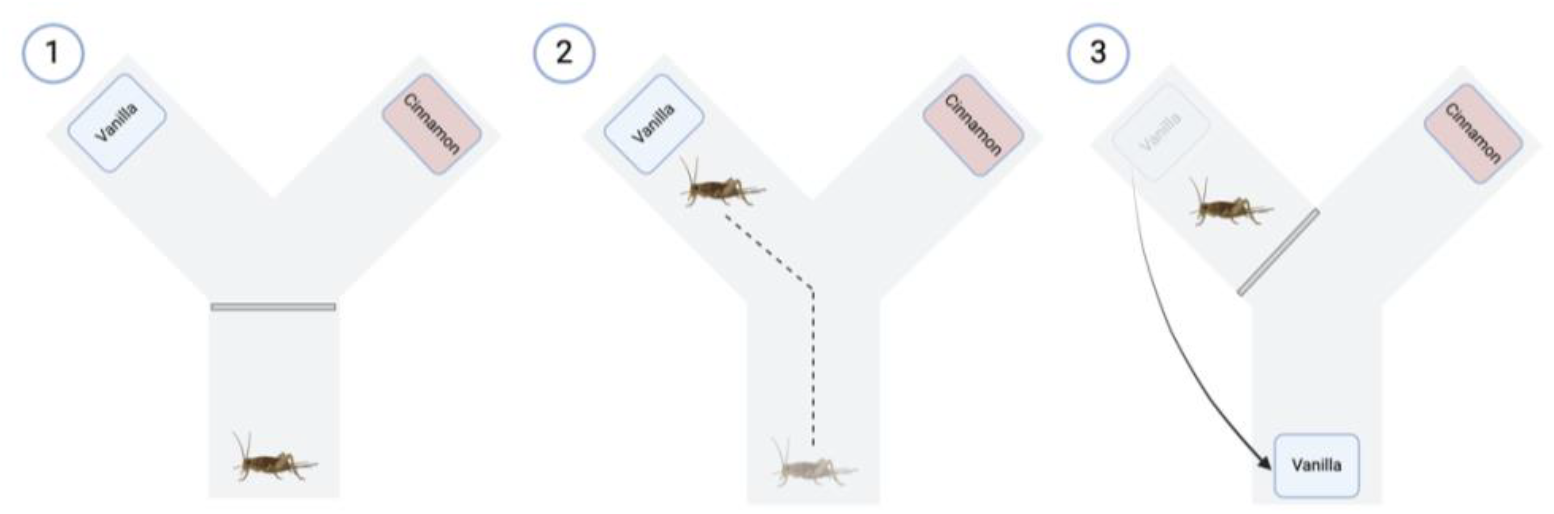
Schematic of scent preference test. Created using Biorender.com.

### Escape learning paradigm

Age-dependent changes in cognitive flexibility were assessed using a modified olfactory escape paradigm based on Scotto-Lomassese et al. (2003), adapted to the ethological and motivational profile of house crickets. The task required individuals to override innate sensory preferences to access a biologically relevant reward, providing a robust assay of learning and decision-making.

### Reinforcement design and ethical considerations

To maximize ecological validity while minimizing stress, rewards consisted of access to a thermally and socially enriched environment containing a heating pad (29 ± 1°C) and acoustic playback of live conspecific calls, both of which elicit strong approach behavior and enhance locomotor activity [31-32]. Incorrect choices led to placement in a sensory-deprived, thermally neutral, and acoustically silent isolation chamber, leveraging the species’ aversion to social isolation [33] without excessive distress. All descent points into chambers were cushioned with sterile cotton to prevent injury.

### Arena setup and sensory manipulation

The apparatus comprised a circular light-controlled arena with two symmetrically placed escape holes (Figure 3). Each hole was associated with an odor cue: vanilla (attractive) or cinnamon (aversive), delivered via filter paper (1% extract diluted 1:1 in distilled water) positioned 7 cm behind the openings to generate a detectable but diffuse scent gradient. To introduce a cognitive conflict, the reinforcement contingency was reversed from natural preference: cinnamon led to the enriched reward chamber, whereas vanilla led to the isolation chamber. This design required suppression of instinctive olfactory biases and formation of novel odor-reward associations [20]. The arena was rotated by 45° between trials to eliminate fixed visual cues, and testing occurred in a dark enclosure under directional overhead lighting to standardize illumination and encourage exploration away from the center.

**Figure 3.**
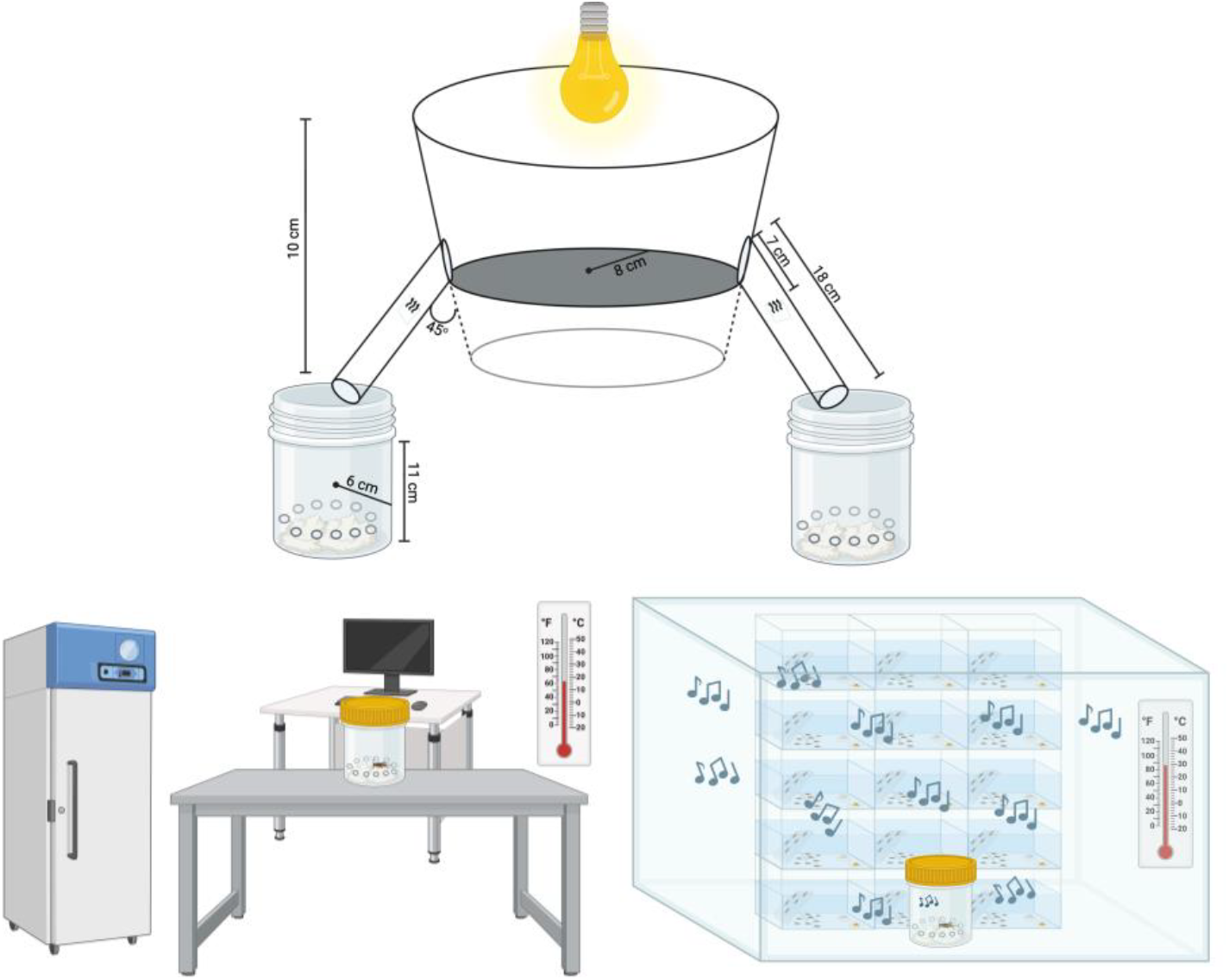
Schematic diagram of the escape paradigm (top). Schematic representations of the positive (right) and the negative (left) reinforcements. Figures created using BioRender.com.

### Trial procedure and behavioral metrics

To increase task engagement without inducing physiological stress, crickets were food-deprived for 48 h prior to testing [20]. Each subject completed 10 trials/day for up to 5 consecutive days. The first two trials were single-choice exposures (only one escape hole open) to ensure experience with both reward and punishment contingencies. The remaining eight trials required free choice between both escape holes. A choice was scored when the cricket’s entire body entered an escape hole, after which it was confined to the outcome chamber for 6 min before the next trial.

Primary measures included: (1) escape hole selection (rewarded vs. punished), (2) decision latency (release-to-choice time), and (3) task acquisition, with “learners” defined as individuals achieving ≥7 correct choices in any 8-trial block. Learners were removed from subsequent sessions; non-learners were those failing to reach criterion after five days.

### Euthanasia and morphometric measurement

Crickets were euthanized in accordance with AVMA guidelines via controlled CO2 exposure, followed by decapitation to ensure death [34]. Morphometric measurements were obtained immediately post-mortem under standardized conditions using digital calipers and acrylic rulers. Body length was measured from the frons to the posterior abdominal tip, excluding cerci, wings, and ovipositor. Antennal length was recorded as the mean of left and right antennae, and hind leg length was measured with the limb fully extended along its natural axis. Specimens with damaged or missing antennae at euthanasia were excluded from antennal analyses. Individuals that died prematurely were excluded from all morphological analyses due to uncertainty in post-mortem interval and the potential effects of tissue degradation or cannibalism.

### Statistical Analysis

Data were stratified by age group and sex before analysis. Binary outcomes from the escape learning paradigm were expressed as n (%) and compared using Pearson’s *χ*^2^ or Fisher’s exact tests; zero counts were adjusted with the modified Haldane-Anscombe correction [35] and effect sizes reported as relative risks (RR, 95% CI). Continuous outcomes (olfactory preference, decision latency) were summarized as mean ± SD and analyzed by one-way or two-way analysis of variance (ANOVA) after Shapiro-Wilk test normality testing, with Tukey’s or Bonferroni-adjusted post hoc tests as appropriate. Effect sizes were expressed as Cohen’s *d* (Hedges’ *g*-corrected) with 95% CIs [36-38].

Morphological effects were assessed using two-way ANOVA, followed by linear regression to identify traits associated with performance. Potential confounders were evaluated with analysis of covariance (ANCOVA). For binary success outcomes, logistic regression was used; in cases of quasi-complete or complete separation, Firth’s penalized likelihood method was applied using the *logistf* package in R [39-40], with odds ratios (OR, 95% CI) from profile penalized likelihood.

Analyses were performed in GraphPad Prism v10.0.3 (GraphPad Software) and R v4.4.0 (Post Software, PBC); α was set at 0.05.

## Results

### Morphological changes across age and sex

Adults exhibited longer antennae than mid-age crickets, driven by females, with higher antennal-to-body and antennal-to-weight ratios than both older groups (*P*’s < 0.05); males consistently exceeded females in antennal-to-weight ratios (P’s < 0.01). Hindleg length was shorter in adults than in mid-age and geriatric crickets, with hindleg-to-weight ratios highest in adults across sexes and males exceeding females at all ages (*P*’s < 0.05). Femoral volume and cross-sectional area (CSA) peaked at mid-age, with larger values in females (*P*’s < 0.05); surface area-to-volume (SA/V) ratios increased with age and were higher in males at mid-age and geriatric stages (*P*’s < 0.01). Detailed ANOVA outputs are in Appendix 1.

### Age-related decline in olfactory discrimination is partly explained by morphological scaling

In the Y-maze assay, adults showed greater preference for the vanilla-scented arm than both mid-age and geriatric crickets (*d*’s = 0.73 to 0.98, *P*’s < 0.05), with no differences between the latter two groups (Figure 4A). This effect was male-driven (*d*’s = 1.11 to 1.28, *P*’s < 0.05), as females showed no age-related changes (Figure 4B). Full statistical output and effect sizes are in Appendix 2.

**Figure 4.**
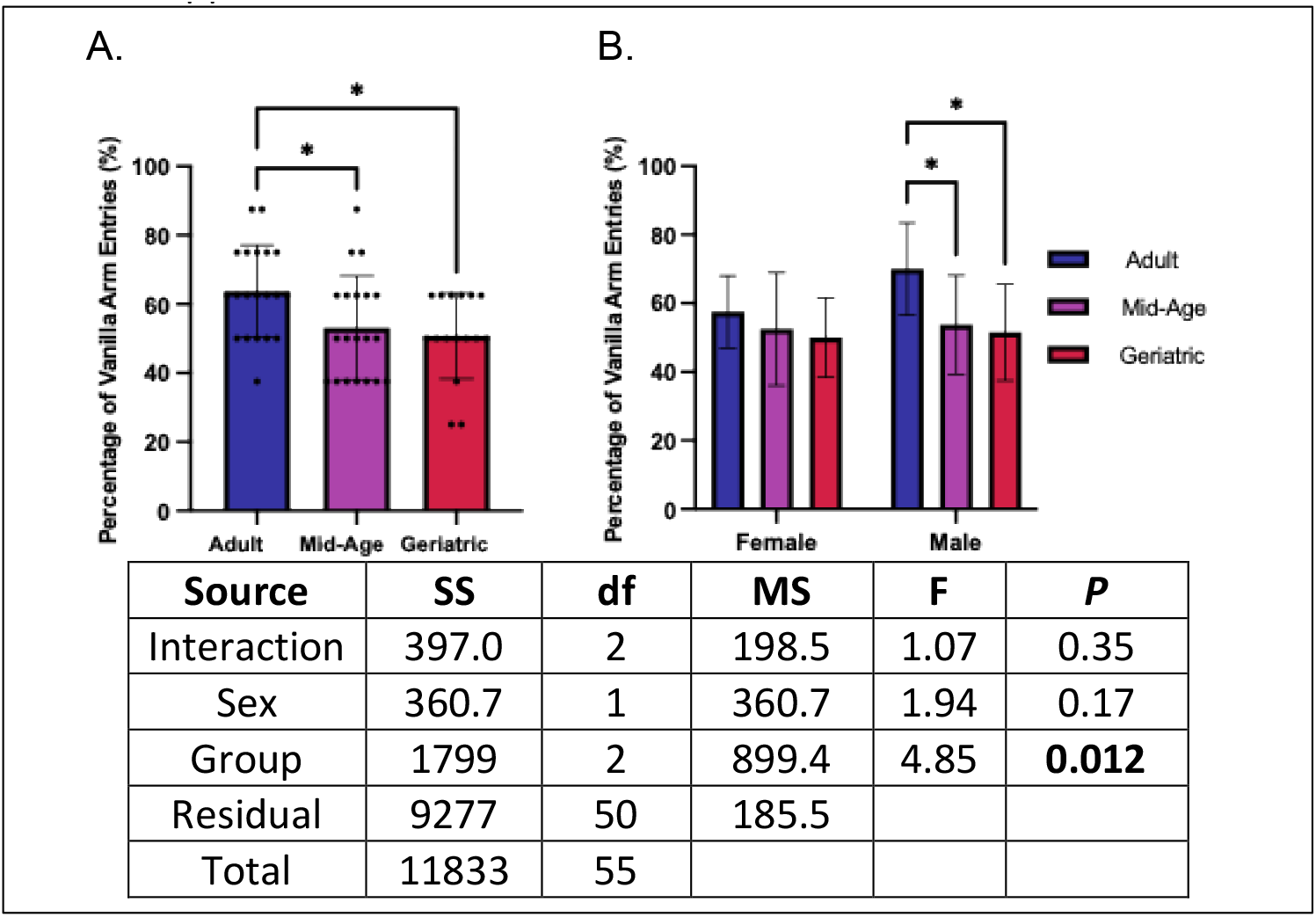
Olfactory recognition memory is sex-dependent and decreases with age. **(A)** Adult crickets had greater vanilla arm preference compared to mid-age and geriatrics. **(B)** Males recapitulated overall trends, while females showed no age-associated changes (\**P* < 0.05).

Morphological predictors of scent preference showed that antennal-to-body length and antennal-to-weight ratios correlated positively with preference, but these associations were attenuated after adjusting for age and sex, eliminating age-group differences. Hindleg-to-weight ratio also correlated with preference but did not persist after adjustment, and other motor or femoral traits showed no relationship. Logistic regression outputs and graphical depictions are in Appendix 3.

### Age-related decline in physiological resilience is linked to femoral morphology

Adults showed greater percent weight gain than mid-age and geriatric crickets (*d*’s = 2.30 to 2.49, *P*’s < 0.0001), with no difference between the latter two groups (Figure 5A). This pattern was evident in both sexes, although males generally retained more mass than females, and mid-age males gained less than geriatrics (Figures 5B-C). Femoral traits were independent predictors of weight change. Larger femoral volume and CSA were associated with reduced mass loss, whereas higher femoral SA/V ratios predicted accelerated decline (Appendix 4 Figure S1, Tables S3-5). These relationships persisted after adjusting for age and sex, indicating that greater muscular investment may buffer against age-associated weight loss.

**Figure 5.**
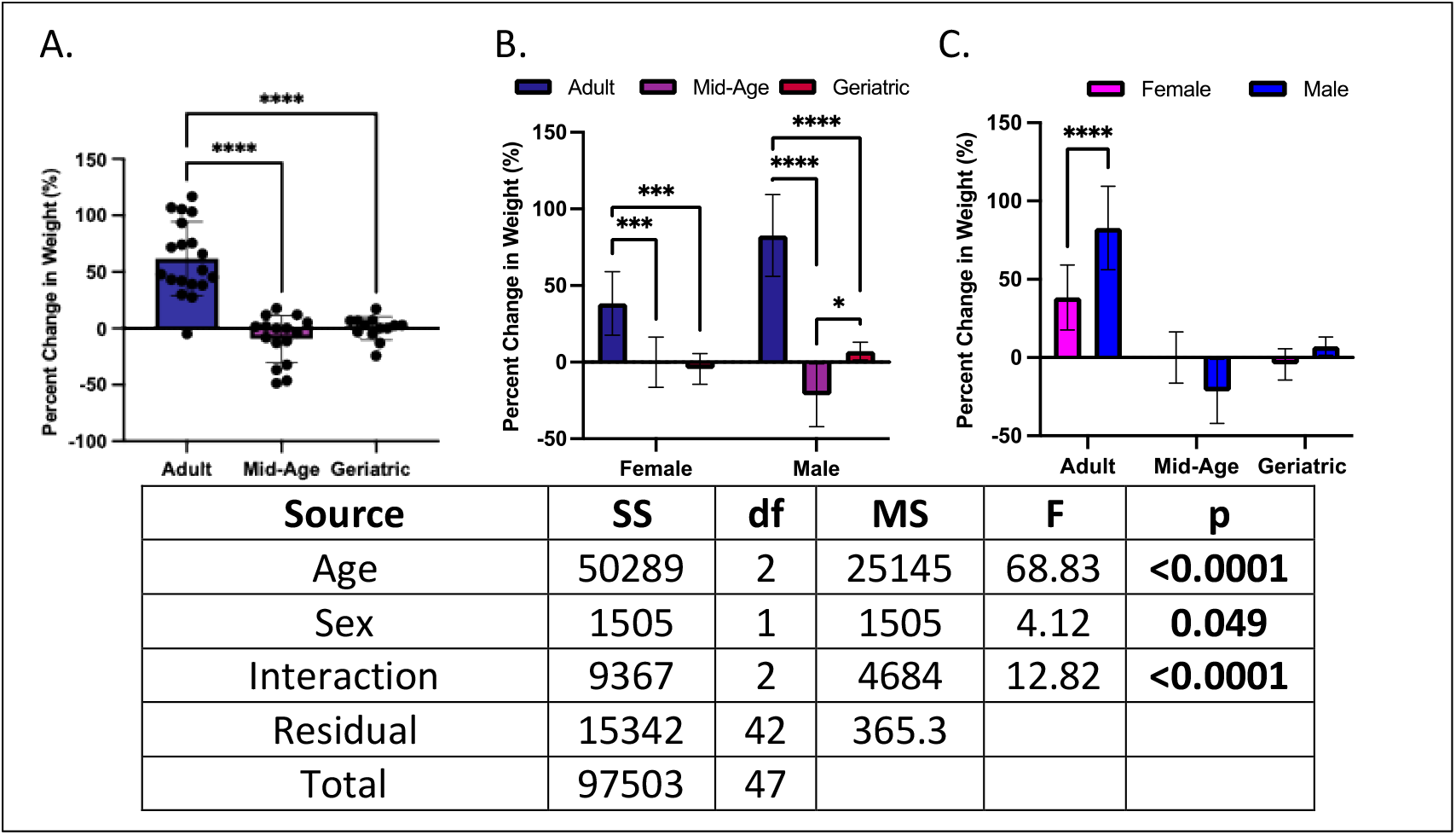
Age- and sex-differences in percent change in weight. **(A)** Adult crickets had higher weight changes than their mid-age and adult counterparts. **(B)** Both sexes generally followed the same trend as the overall population. **(C)** Adult males had higher weight changes than females (\**P* < 0.05, \*\*\**P* < 0.001, \*\*\*\**P* < 0.0001).

### Escape learning performance declines with age, with mid-age crickets showing prolonged latencies independent of morphology

Several crickets were excluded due to mortality before or during testing, including one mid-aged male, one geriatric male, and one geriatric female prior to final weight measurement, and three additional individuals (one mid-aged male, one geriatric male, one geriatric female) during the first two days of testing. All exclusions were incorporated into statistical analyses.

Task acquisition declined with age, with geriatrics performing worse than adults and mid-age crickets across the five-day paradigm (Figure 6A; Appendix 5 Tables S1-2). Early in testing, geriatrics achieved lower success rates than adults (*RR*’s = 2.28 to 7.20, *P*’s = 0.04 to 0.0004), but not mid-age crickets; by days four and five, both younger groups consistently outperformed geriatrics (*RR*’s = 1.70 to 2.00, *P*’s = 0.04 to 0.001). Age effects were stronger in males, with adults outperforming geriatrics from day 2 onward (*RR*’s = 2.22 to 13.5, *P*’s < 0.05), whereas female differences were transient, with adults showing an advantage over geriatrics on day 3 only (*RR* = 2.22 [95% CI: 0.93, 5.28], *P* = 0.029) (Figure 6B-C). Adjustment for sensory morphological traits indicated that only antennal-to-weight ratio attenuated early adult-geriatrics differences (*P*’s > 0.05); from day 3 onward, age effects persisted (*P*’s < 0.01), suggesting true cognitive decline rather than morphological limitation.

**Figure 6.**
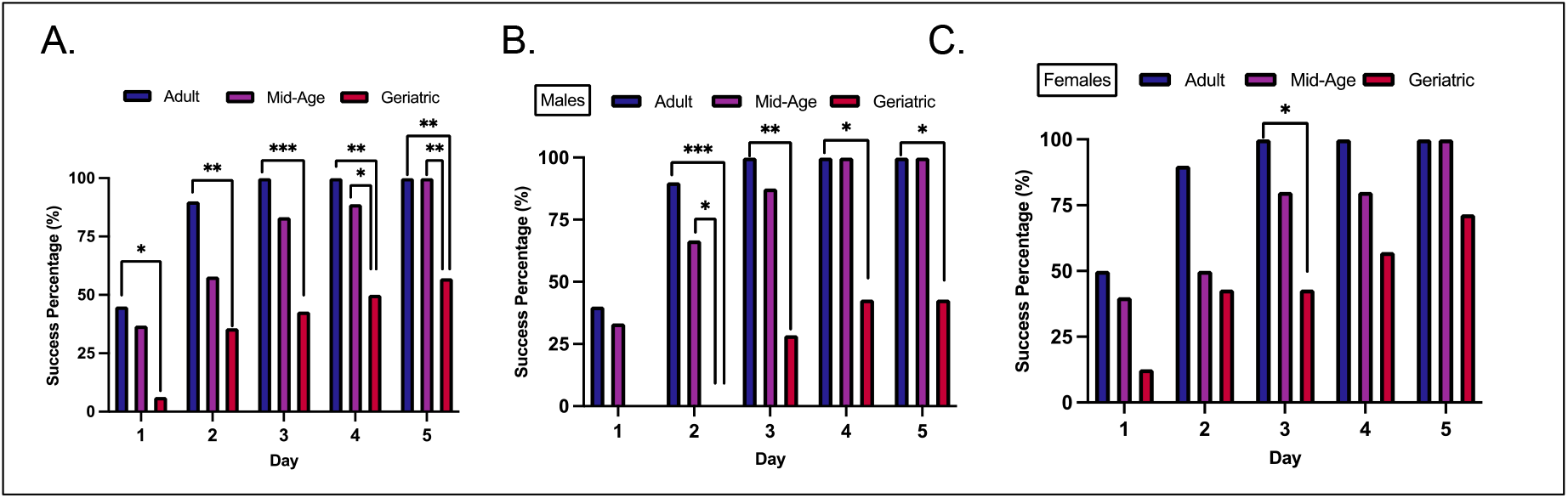
Age- and sex-differences in success percentages. **(A)** Geriatric crickets performed worse on learning tasks than their mid-age and adult counterparts. **(B)** Geriatric males followed similar trends as the overall population, while **(C)** differences among female cohorts were less pronounced (\**P* < 0.05, \*\**P* < 0.01, \*\*\**P* < 0.001).

Across training and test phases, mid-aged crickets consistently showed longer decision-making times than adults or geriatrics (*d*’s = −1.53 to 1.05, *P*’s < 0.05), with the effect evident in both sexes (Figure 7; Appendix 5 Tables S3-4). Several sensory and motor traits, including antennal investment and hindleg or femoral dimensions, correlated with decision time in univariate models, but none explained age effects after adjustment, indicating morphology did not account for prolonged mid-age latencies (Appendix 6 Figures S1-2, Tables S1-10).

**Figure 7.**
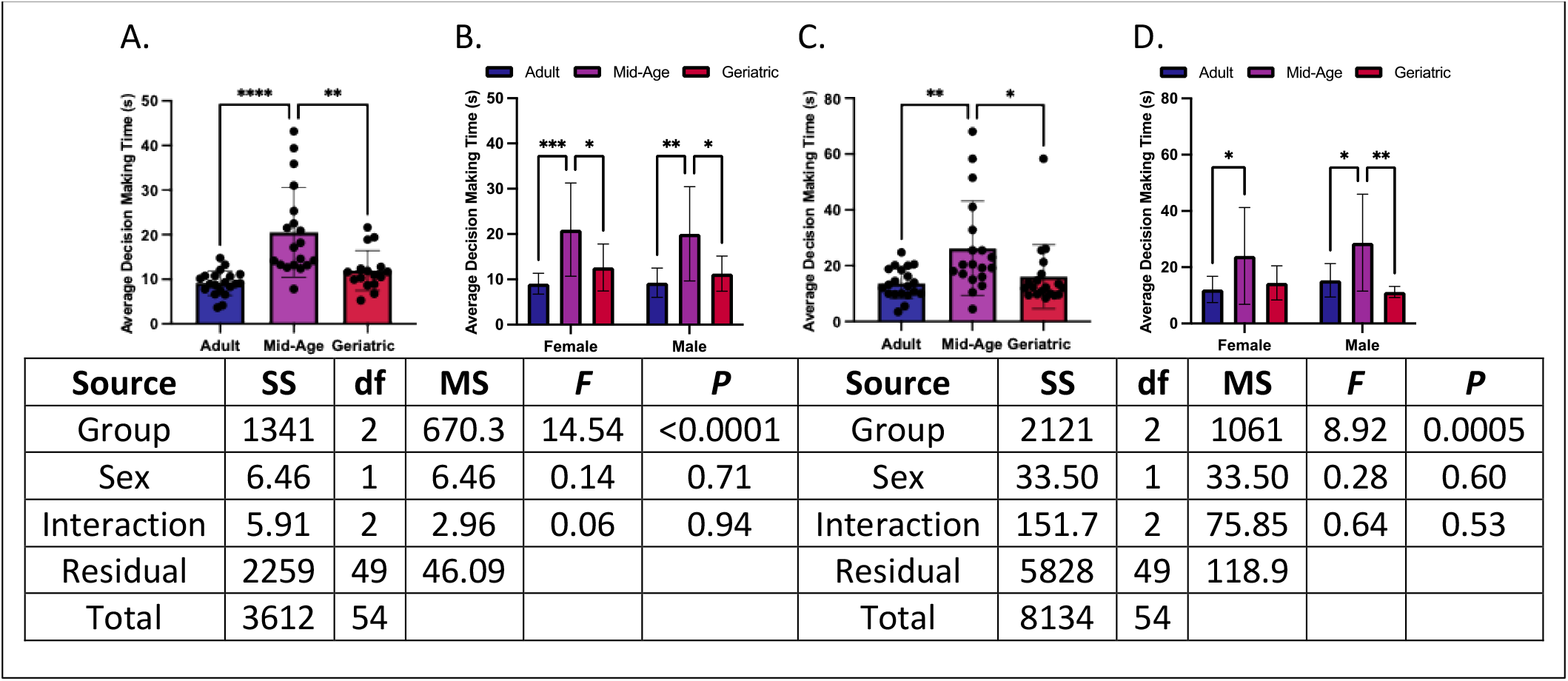
Age- and sex-related differences in decision-making time. **(A-B)** During main trials, mid-age crickets displayed longer average decision-making times compared to adult and geriatric cohorts, with consistent patterns in both females and males. **(C-D)** This trend persisted during test trials, where mid-age crickets again required more time to decide compared to both adult and geriatrics; effects were robust across sexes (\**P* < 0.05, \*\**P* < 0.01, \*\*\*\**P* < 0.0001).

In the learning phase, mid-age crickets also took longer to reach the punishment arm than adults or geriatrics (*d*’s = −1.11 to 0.85, *P*’s < 0.05) (Figures 8A-B), an effect absent in the test phase (Figure 8C-D). Femoral SA/V ratio partially attenuated this difference, whereas other traits had no effect (Appendix 6 Figures S3-4, Tables S11-12). Time to reward arm was consistently longer for mid-age crickets than for adults in both learning and test phases (*d*’s = −1.42 to 1.64, *p*’s < 0.05) (Figure 8E-H), with no consistent sex differences. Morphological traits did not explain these delays in adjusted analyses, indicating that age group remained the primary determinant of reward acquisition time (Appendix 6 Figure S5-6, Tables S13-22).

**Figure 8.**
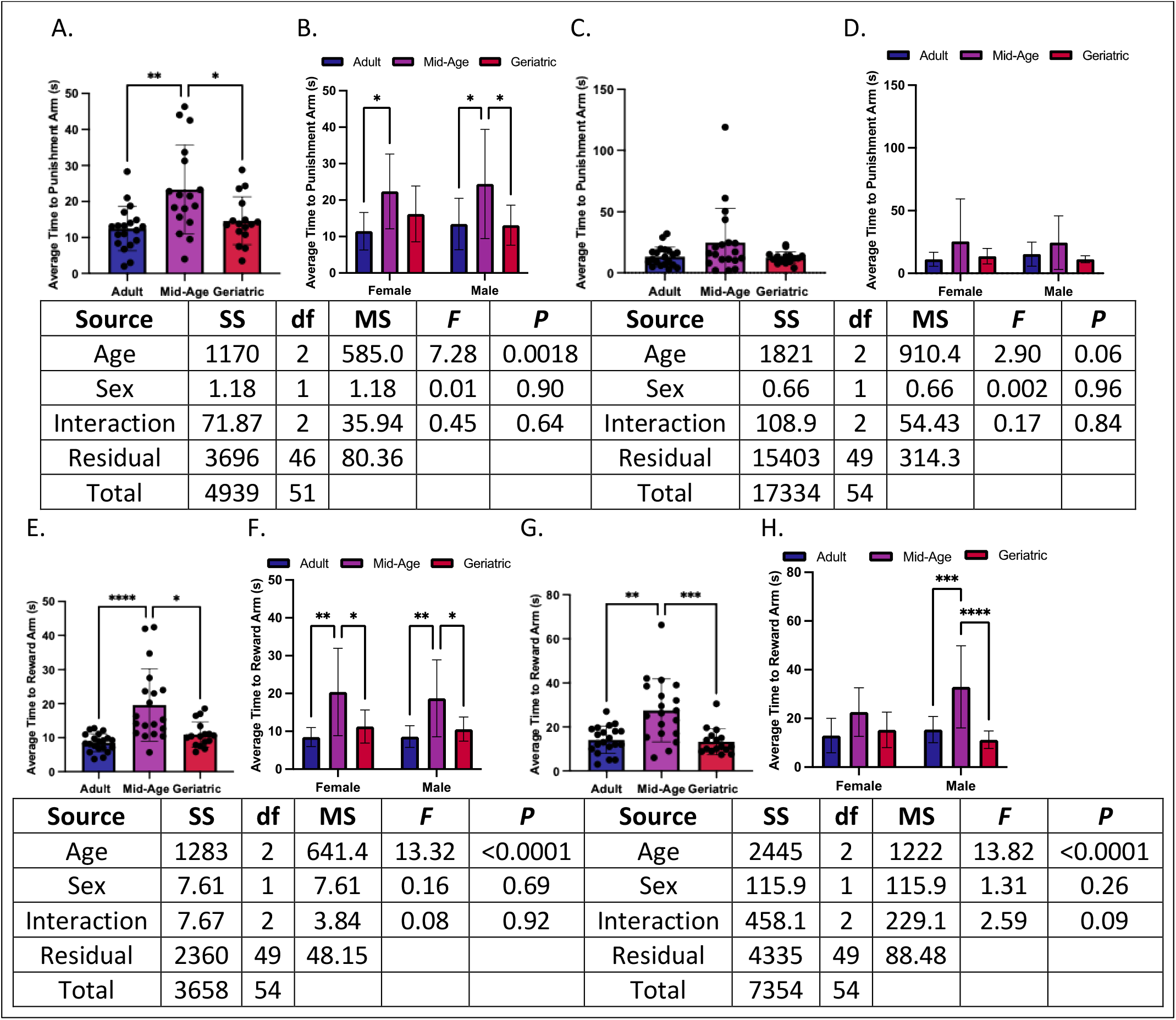
Age- and sex-related differences in average time to choice arms. **(A-B)** Time to reach the punishment arm was elevated in mid-age crickets during main trials, relative to both adults and geriatrics, with consistent delays in females and males. **(C-D)** No age- or sex-based differences emerged for punishment arm times during test trials. **(E-F)** Mid-age crickets showed prolonged times to reach the reward arm during main trials relative to other age groups; these differences were conserved across sexes. **(G-H)** In test trials, both female and male mid-age crickets again demonstrated longer reward arm times compared to adults and geriatrics, particularly in males (\**P* < 0.05, \*\**P* < 0.01, \*\*\**P* < 0.001, \*\*\*\**P* < 0.0001).

## Discussion

Our study demonstrates that house crickets exhibit age-related cognitive decline characterized by impairments in sensory discrimination, associative learning, decision-making speed, and physiological resilience. Using a dual-task design (i.e., spontaneous olfactory preference and escape-based learning), we establish this invertebrate as a scalable model for studying conserved mechanisms of neural aging. Geriatric crickets showed reduced olfactory preference and impaired learning, while mid-aged individuals exhibited delayed decision-making, indicating that decline begins well before late life [41]. This trajectory mirrors that of vertebrates, where cognitive function peaks in early adulthood and declines thereafter [17], highlighting fundamental, phylum-spanning processes underlying neural aging [7].

A central question in aging research is whether cognitive decline stems primarily from peripheral deterioration or intrinsic neural aging. We observed age-related morphological changes in crickets (i.e., shorter antennae, altered limb proportions, weight loss) that parallel human declines in sensory and musculoskeletal integrity [42]. Crickets with larger antennae performed better in odor discrimination, and those with larger hindlegs exhibited less weight loss, suggesting that preserved peripheral structure can buffer certain aging effects. However, adjusting for these traits did not eliminate cognitive impairments in geriatric crickets. Even with intact morphology, older individuals remained deficient in learning and memory, pointing to age-related changes in central neural circuits [43]. This aligns with vertebrate models, where cognitive decline reflects synaptic dysfunction and altered neural plasticity rather than peripheral deficits alone [44].

Olfactory behavior provides a particularly sensitive readout of this central decline, given its reliance on both peripheral sensory input and higher-order processing. Olfactory decline emerged as a prominent feature of aging in male crickets, which progressively lost their innate preference for vanilla odor in the Y-maze. This mirrors findings in humans, where olfactory impairment is an early and predictive marker of cognitive decline and neurodegeneration [45]. The conservation of this relationship suggests that olfaction may serve as a cross-species indicator of neural aging. Interestingly, female crickets maintained stable odor preference with age, revealing a sex-specific divergence. Similar sex differences are observed in humans, where males typically show greater olfactory and cognitive decline, potentially due to hormonal or neurobiological factors [46-47]. The resilience observed in female crickets may reflect enhanced sensory maintenance or plasticity, warranting further study into sex-specific mechanisms such as neurogenesis or neuromodulator regulation in aging brains [48-49].

Building on the observed sensory decline, age-related impairments in associative learning further underscore the breadth of cognitive deterioration in older crickets. In the escape task, which required forming an odor-reward association, geriatric individuals struggled to acquire the correct response pattern, often failing to reach learning criteria despite repeated training. Younger adults quickly learned to escape via the rewarded odor cue, whereas older crickets exhibited persistent errors, indicative of deficits in memory formation, recall, or behavioral flexibility [50]. Given the central role of the mushroom bodies in insect learning, age-related dysfunction in these structures likely contributes to the cognitive decline we observed [51]. Prior work shows that adult-born neurons continue to populate mushroom bodies throughout life, but this process diminishes with age [11], paralleling reduced plasticity in the aging mammalian brain [52]. Importantly, the observed learning impairments persisted even when motor ability and motivation were controlled, pointing to a genuine decline in central cognitive function rather than peripheral limitations.

Notably, cognitive aging in crickets did not follow a strictly linear trajectory. While mid-aged individuals (8 weeks old) exhibited the slowest decision-making in the escape task, geriatrics retained faster but less accurate response strategies. Aging in this case, may involve shifts in behavioral strategy, where older crickets may favor impulsive or phototactically driven responses, while mid-aged individuals may adopt a more cautious, deliberative approach that prolongs decision time. Such trade-offs between speed and accuracy are well-documented in human cognitive aging and may reflect changes in motivation, risk aversion, or strategy selection with age [53]. These findings highlight the complexity of aging trajectories and underscore the need to consider non-linear and individual-specific dynamics in cognitive decline.

Our findings also reveal a decline in physiological resilience with age, paralleling patterns of frailty observed in mammals. Across the experimental period, adult crickets maintained or gained weight, whereas mid-aged and geriatric cohorts experienced progressive weight loss, despite identical nutritional access. In humans, unintentional weight loss in late life is a hallmark of frailty and diminished stress tolerance, often signaling systemic decline under physiological challenge [54]. The behavioral assays, comprising repeated handling and mild aversive stimuli, appeared to impose a greater metabolic burden on older crickets. Notably, femoral morphology emerged as a predictor of resilience: geriatric crickets with greater femur volume and cross-sectional area lost less weight, suggesting that preserved muscle mass conferred resistance to stress-induced wasting. This parallels human aging, where sarcopenia is closely linked to frailty, impaired recovery, and elevated mortality risk [54]. Beyond mobility, skeletal muscle serves as a metabolic buffer, supporting energy demands during stress [55]. Our data suggest that, as in humans, muscle condition in crickets modulates vulnerability in late life, a finding that invites further exploration into whether interventions targeting musculoskeletal integrity can enhance survival or cognitive function in aging invertebrates.

While this study provides key insights into behavioral aging in an invertebrate model, several limitations warrant consideration. First, although cohort sizes were adequate to detect large effects, sample sizes for sex-stratified and morphology-adjusted analyses were modest, leading to wide confidence intervals for some estimates. Larger cohorts across age and sex will be necessary to validate observed trends and support interaction modeling. Second, although we identified morphological correlates of behavior and resilience, we did not directly assess underlying cellular or molecular pathology. As the absence of geropathological data limits our ability to attribute cognitive decline to specific neurobiological changes, we are currently developing the first validated geropathology grading platform for crickets, including the presence of neuronal lipofuscin in the brain as well as various other hallmarks of aging throughout the various organ systems. Future work combining histopathology with behavioral phenotyping will be critical for disentangling structural and functional contributors to cognitive decline and for identifying conserved hallmarks of aging across species.

## Supporting information

BioRender Publication Licenses

