## Supplemental File for "Age-related cognitive decline in house crickets reveals conserved patterns of sensory and learning deficits across the lifespan"

Appendix 1. Morphological Changes Across Age and Sex

***Sensory Morphology***

A.	B.

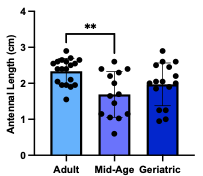

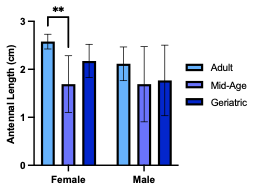

|  | **SS** | **df** | **MS** | **F** | ***P*** |
| --- | --- | --- | --- | --- | --- |
| Interaction | 0.45 | 2 | 0.23 | 0.87 | 0.43 |
| Sex | 0.98 | 1 | 0.98 | 3.78 | 0.058 |
| Group | 3.42 | 2 | 1.71 | 6.59 | **0.0032** |
| Residual | 11.16 | 43 | 0.26 |  |  |
| Total | 16.01 | 48 |  |  |  |

**Figure S1. Antennal length as a function of age and sex.** **(A)** Adults displayed greater antennal length compared to mid-aged crickets. **(B)** Sex-stratified analysis showed that this difference was restricted to females, with adult females exhibiting longer antennae than mid-aged females, while males showed no age-related variation. Values represent mean ± SD (***P* < 0.01).

A.	B.

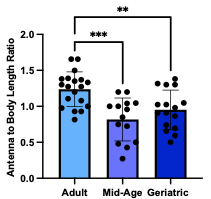

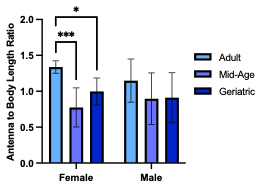

|  | **SS** | **df** | **MS** | **F** | ***P*** |
| --- | --- | --- | --- | --- | --- |
| Interaction | 0.19 | 2 | 0.09 | 1.30 | 0.28 |
| Sex | 0.03 | 1 | 0.03 | 0.44 | 0.51 |
| Group | 1.44 | 2 | 0.72 | 10.06 | **0.0003** |
| Residual | 3.08 | 43 | 0.07 |  |  |
| Total | 4.74 | 48 |  |  |  |

**Figure S2. Age- and sex-specific differences in antennal investment relative to body length.** **(A)** Adults exhibited a higher antennal-to-body length ratio than mid-aged crickets. **(B)** This pattern was driven by females, with adult females showing an elevated antennal-to-body length ratio compared to mid-aged females, whereas males exhibited no age-related differences. Data are presented as mean ± SD (**P* < 0.05, ***P* < 0.01, ****P* < 0.001).

A.	B.	C.

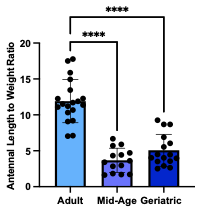

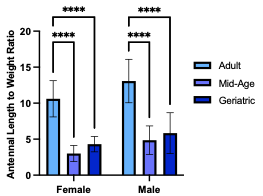

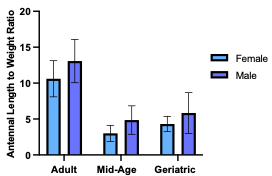

|  | **SS** | **df** | **MS** | **F** | ***P*** |
| --- | --- | --- | --- | --- | --- |
| Interaction | 1.96 | 2 | 0.98 | 0.19 | 0.83 |
| Sex | 44.35 | 1 | 44.35 | 8.56 | **0.0055** |
| Group | 614.9 | 2 | 307.5 | 59.33 | **<0.0001** |
| Residual | 222.8 | 43 | 5.18 |  |  |
| Total | 884.0 | 48 |  |  |  |

**Figure S3. Antennal length relative to body weight varies with age but not sex. (A)** Adults displayed a higher antennal-to-weight ratio than their older counterparts. **(B)** This age-related pattern was observed in both females and males, each showing elevated ratios in adulthood compared to mid-age and geriatrics. **(C)** Within each age group, antennal-to-weight ratios did not differ between sexes. Data are shown as mean ± SD (**P* < 0.05, ***P* < 0.01, ****P* < 0.001).

***Motor Morphology***

A.	B.	C.

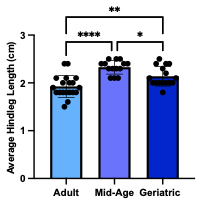

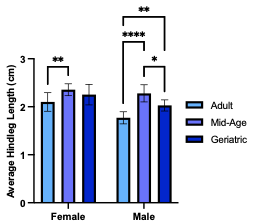

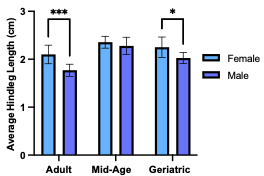

|  | **SS** | **df** | **MS** | **F** | ***P*** |
| --- | --- | --- | --- | --- | --- |
| Interaction | 0.12 | 2 | 0.06 | 2.41 | 0.10 |
| Sex | 0.51 | 1 | 0.51 | 20.01 | **<0.0001** |
| Group | 1.15 | 2 | 0.57 | 22.28 | **<0.0001** |
| Residual | 1.11 | 43 | 0.03 |  |  |
| Total | 2.89 | 48 |  |  |  |

**Figure S4. Age- and sex-dependent variation in hindleg length.** **(A)** Hindleg length peaked in mid-aged crickets, exceeding that of both adults and geriatrics, with geriatrics also exhibiting longer hindlegs than adults. **(B)** Males mirrored the overall pattern, whereas in females, mid-aged individuals only surpassed adults. **(C)** Sex differences were evident only in adult and geriatric cohorts, where females possessed longer hindlegs than males. Data represent mean ± SD (**P* < 0.05, ***P* < 0.01, ****P* < 0.001, *****P* < 0.0001).

A.	B.

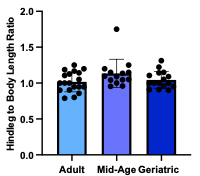

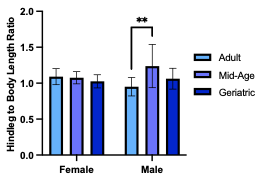

|  | **SS** | **df** | **MS** | **F** | ***P*** |
| --- | --- | --- | --- | --- | --- |
| Interaction | 0.18 | 2 | 0.091 | 4.49 | **0.017** |
| Sex | 0.01 | 1 | 0.01 | 0.22 | 0.64 |
| Group | 0.15 | 2 | 0.08 | 3.77 | **0.031** |
| Residual | 0.88 | 43 | 0.02 |  |  |
| Total | 1.22 | 48 |  |  |  |

**Figure S5. Hindleg investment relative to body length across age and sex. (A)** No overall differences in hindleg-to-body length ratio were observed among age groups. **(B)** Sex-stratified analysis revealed that mid-age males exhibited higher ratios than adult males, whereas no differences were detected among females. Data are presented as mean ± SD (***P* < 0.01).

A.	B.	C.

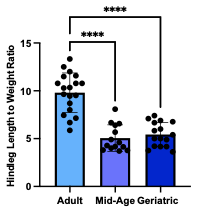

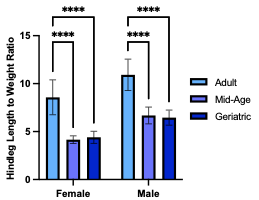

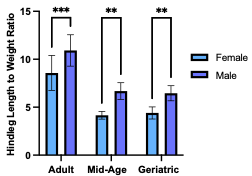

|  | **SS** | **df** | **MS** | **F** | ***P*** |
| --- | --- | --- | --- | --- | --- |
| Interaction | 0.41 | 2 | 0.21 | 0.14 | 0.87 |
| Sex | 62.17 | 1 | 62.17 | 43.13 | **<0.0001** |
| Group | 213.6 | 2 | 106.8 | 74.08 | **<0.0001** |
| Residual | 61.98 | 43 | 1.44 |  |  |
| Total | 338.16 | 48 |  |  |  |

**Figure S6. Age- and sex-related differences in hindleg length relative to body weight. (A)** Adults exhibited higher hindleg-to-weight ratios than mid-age and geriatrics. **(B)** This age-related trend was consistent in both sexes, with adult males and females each showing elevated ratios compared to their older counterparts. **(C)** Across all age groups, males maintained higher hindleg-to-weight ratios than females. Data are presented as mean ± SD (***P* < 0.01, ****P* < 0.001, *****P* < 0.0001).

***Femur Morphology***

A.	B.	C.

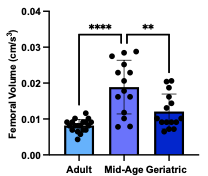

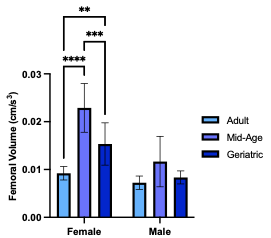

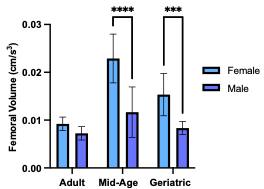

|  | **SS** | **df** | **MS** | **F** | ***P*** |
| --- | --- | --- | --- | --- | --- |
| Interaction | 1.68e-04 | 2 | 8.41e-05 | 7.05 | **0.0023** |
| Sex | 5.17e-04 | 1 | 5.17e-04 | 43.29 | **<0.0001** |
| Group | 6.25e-04 | 2 | 3.12e-04 | 26.18 | **<0.0001** |
| Residual | 5.01e-04 | 42 | 1.19e-05 |  |  |
| Total | 1.81e-03 | 47 |  |  |  |

**Figure S7. Age- and sex-specific variation in femoral volume. (A)** Femoral volume was greatest in mid-aged crickets, exceeding that of both adult and geriatric cohorts. **(B)** Among females, mid-aged individuals exhibited the largest femoral volume, followed by geriatrics and then adults, whereas males showed no differences across ages. (C) Mid-aged and geriatric females possessed larger femoral volumes than their male counterparts. Data are presented as mean ± SD (***P* < 0.01, ****P* < 0.001, *****P* < 0.0001).

A.	B.	C.

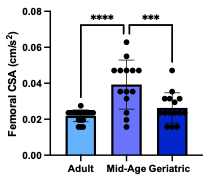

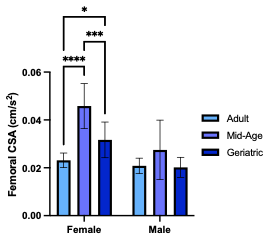

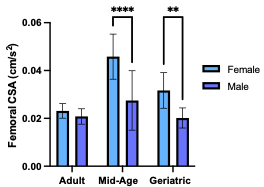

|  | **SS** | **df** | **MS** | **F** | ***P*** |
| --- | --- | --- | --- | --- | --- |
| Interaction | 5.09e-04 | 2 | 2.55e-04 | 5.37 | **0.0084** |
| Sex | 1.31e-03 | 1 | 1.31e-03 | 27.53 | **<0.0001** |
| Group | 1.69e-03 | 2 | 8.46e-04 | 17.84 | **<0.0001** |
| Residual | 1.99e-03 | 42 | 4.74e-05 |  |  |
| Total | 5.50e-03 | 47 |  |  |  |

**Figure S8. Age- and sex-dependent differences in femoral cross-sectional area (CSA). (A)** Femoral CSA was greatest in mid-age crickets, surpassing values observed in both adults and geriatrics. **(B)** Among females, mid-aged individuals exhibited the largest CSA, followed by geriatrics and then adults, whereas males showed no age-related differences. (C) Mid-aged and geriatric females had larger femoral CSA than males within the same age groups. Data are shown as mean ± SD (***P* < 0.01, ****P* < 0.001, *****P* < 0.0001).

A.	B.	C.

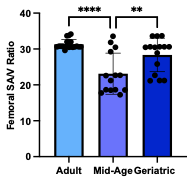

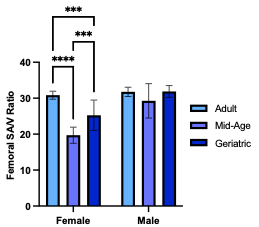

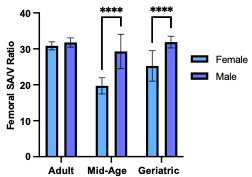

|  | **SS** | **df** | **MS** | **F** | ***P*** |
| --- | --- | --- | --- | --- | --- |
| Interaction | 155.5 | 2 | 77.74 | 10.93 | **0.0002** |
| Sex | 371.7 | 1 | 371.7 | 52.26 | **<0.0001** |
| Group | 356.0 | 2 | 178.0 | 25.03 | **<0.0001** |
| Residual | 298.8 | 42 | 7.113 |  |  |
| Total | 1182 | 47 |  |  |  |

**Figure S9. Age- and sex-related variation in femoral surface area-to-volume (SA/V) ratio. (A)** Mid-aged crickets exhibited the lowest femoral SA/V. (B) Among females, SA/V was lowest in mid-aged individuals, followed by geriatrics and then adults, whereas males showed no age-dependent differences. (C) Mid-aged and geriatric females displayed smaller femoral SA/V ratios than males of the same age. Data represent mean ± SD (***P* < 0.01, ****P* < 0.001, *****P* < 0.0001).

**Appendix 2. Olfactory discrimination declines with age.**

|  | Group [Mean (SD)] | | | Adult-Mid-Age | | Adult-Geriatric | | Mid-Age-Geriatric | |
| --- | --- | --- | --- | --- | --- | --- | --- | --- | --- |
|  | Adult | Mid-Age | Geriatric | *d*  (95% CI) | Adj. *P-*Value | *d*  (95% CI) | Adj. *P-*Value | *d*  (95% CI) | Adj. *P-*Value |
| Overall | N = 20 | N = 20 | N = 16 | 0.73  (0.09, 1.37) | **0.047** | 0.98  (0.28, 1.67) | **0.019** | 0.16  (-0.49, 0.82) | 0.87 |
|  | 63.75 (13.39) | 53.13 (15.11) | 50.78 (12.47) |  |  |  |  |  |  |
| Female | N = 10 | N = 10 | N = 8 | 0.35  (-0.54, 1.23) | 0.69 | 0.65  (-0.30, 1.60) | 0.48 | 0.16  (-0.77, 1.10) | 0.92 |
|  | 57.50 (10.54) | 52.50 (16.46) | 50.00 (11.57) |  |  |  |  |  |  |
| Male | N = 10 | N = 10 | N = 8 | 1.11  (0.17, 2.06) | **0.027** | 1.28  (0.26, 2.30) | **0.017** | 0.15  (-0.79, 1.08) | 0.94 |
|  | 70.00 (13.44) | 53.75 (14.49) | 51.56 (14.07) |  |  |  |  |  |  |

**Table S1. Group means and effect sizes for scent preference test across age groups.** Mean values with standard deviations (SD) are reported for percentage of vanilla arm entries in adult, mid-age, and geriatric crickets (N = number of individuals per group). Pairwise comparisons are quantified using Cohen’s *d* with Hedges’ *g* correction to account for small sample size bias. Effect sizes are reported alongside 95% confidence intervals (CI) in the format (lower, upper). Adults served as the reference group for adult-mid-age and adult-geriatric comparisons, while mid-age served as the reference group for mid-age-geriatric comparisons. Adjusted *P*-values were calculated using Tukey’s Honestly Significant Difference (HSD) post-hoc test.

**Appendix 3. Morphological predictions of scent preference performance.**

A.	B.	C.

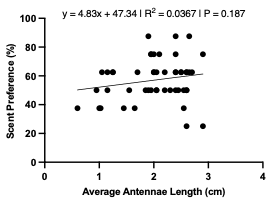

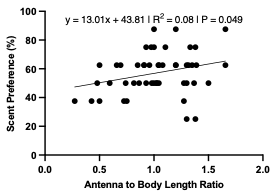

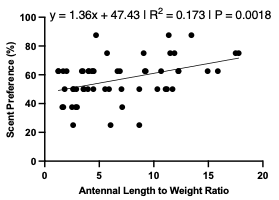

D.	E.	F.

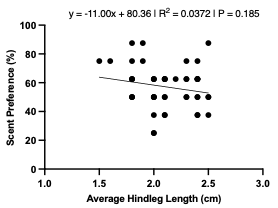

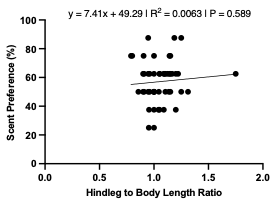

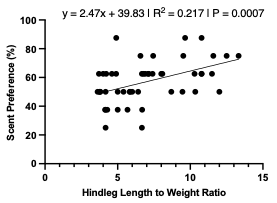

G.	H.	I.

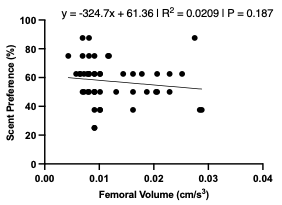

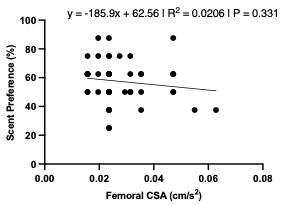

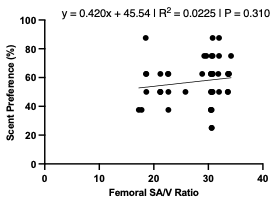

**Figure S1. Relationships between morphological traits and scent preference performance. (A)** Average antennae length showed no association with scent preference. **(B)** Antennal-to-body length ratio positively predicted scent preference, indicating that proportionally longer antennae relative to body size enhanced olfactory performance. **(C)** Antennal-to-weight ratio was also positively associated with scent preference, suggesting that lighter individuals with longer antennae demonstrated superior task success. **(D)** Hindleg length did not correlate with scent preference. **(E)** Hindleg-to-body length ratio likewise showed no relationship with performance. **(F)** Hindleg-to-weight ratio was positively associated with scent preference, reflecting a potential link between proportional hindleg investment and behavioral outcomes. **(G-I)** Femoral volume, cross sectional area (CSA), and surface area-to-volume (SA/V) ratio each showed no association with scent preference.

| **Predictor** | **Estimate** | **Standard error** | **95% CI** | ***P*** |
| --- | --- | --- | --- | --- |
| Intercept | 48.80 | 10.21 | 28.23 to 69.38 | **<0.0001** |
| Group[Mid-Age] | -3.43 | 5.77 | -15.07 to 8.20 | 0.56 |
| Group[Geriatric] | -10.26 | 5.05 | -20.44 to -0.09 | **0.048** |
| Sex[M] | 7.25 | 3.93 | -0.68 to 15.17 | 0.072 |
| Antenna to Body Length | 9.05 | 7.46 | -5.98 to 24.08 | 0.23 |
| Model R^2^ = 0.226. P values adjusted via Holm-Bonferroni correction. Reference group = Adult, Female. | | | | |

**Table S1. Logistic regression evaluating the effects of age, sex, and antennal investment relative to body length on task success in the scent preference assay.**

| **Predictor** | **Estimate** | **Standard error** | **95% CI** | ***P*** |
| --- | --- | --- | --- | --- |
| Intercept | 44.88 | 10.32 | 24.09 to 65.68 | **<0.0001** |
| Group[Mid-Age] | 3.90 | 8.46 | -13.15 to 20.96 | 0.65 |
| Group[Geriatric] | -3.20 | 7.53 | -18.38 to 11.97 | 0.67 |
| Sex[M] | 3.76 | 4.24 | -4.78 to 12.30 | 0.38 |
| Antennal Length to Weight | 1.42 | 0.88 | -0.37 to 3.21 | 0.12 |
| Model R^2^ = 0.226. P values adjusted via Holm-Bonferroni correction. Reference group = Adult, Female. | | | | |

**Table S2. Logistic regression assessing the influence of age, sex, and antennal investment relative to body weight on scent preference task success.**

| **Predictor** | **Estimate** | **Standard error** | **95% CI** | ***P*** |
| --- | --- | --- | --- | --- |
| Intercept | 38.16 | 14.99 | 7.95 to 68.36 | **0.015** |
| Group[Mid-Age] | 3.92 | 8.77 | -13.76 to 21.59 | 0.66 |
| Group[Geriatric] | -1.72 | 8.59 | -19.03 to 15.58 | 0.84 |
| Sex[M] | 0.67 | 5.48 | -10.37 to 11.70 | 0.90 |
| Hindleg Length to Weight | 2.58 | 1.69 | -0.82 to 5.99 | 0.13 |
| Model R^2^ = 0.226. P values adjusted via Holm-Bonferroni correction. Reference group = Adult, Female. | | | | |

**Table S3. Logistic regression examining the effects of age, sex, and hindleg investment relative to body weight on scent preference task success.**

**Appendix 4. Percent weight change and associations with select morphological traits.**

| **Percent Weight Change (%)** | Group [Mean (SD)] | | | Adult-Mid-Age | | Adult-Geriatric | | Mid-Age-Geriatric | |
| --- | --- | --- | --- | --- | --- | --- | --- | --- | --- |
|  | Adult  (N = 19) | Mid-Age  (N = 16) | Geriatric  (N = 13) | *d*  (95% CI) | Adj. *P-*Value | *d*  (95% CI) | Adj. *P-*Value | *d*  (95% CI) | Adj. *P-*Value |
| Overall | 61.72  (32.63) | -9.45  (20.81) | 0.06  (10.23) | 2.49  (1.61, 3.38) | **<0.0001** | 2.30  (1.40, 3.21) | **<0.0001** | -0.55  (-1.29, 0.20) | 0.56 |
|  | Group [Mean (SD)] | | | Adult-Mid-Age | | Adult-Geriatric | | Mid-Age-Geriatric | |
|  | Adult  (N = 9) | Mid-Age  (N = 9) | Geriatric  (N = 8) | *d*  (95% CI) | Adj. *P-*Value | *d*  (95% CI) | Adj. *P-*Value | *d*  (95% CI) | Adj. *P-*Value |
| Female | 38.38  (20.74) | 0.00  (16.35) | -4.45  (9.97) | 1.96  (0.83, 3.08) | **0.0003** | 2.45  (1.19, 3.71) | **0.0001** | 0.31  (-0.65, 1.26) | 0.88 |
|  | Group [Mean (SD)] | | | Adult-Mid-Age | | Adult-Geriatric | | Mid-Age-Geriatric | |
|  | Adult  (N = 10) | Mid-Age  (N = 7) | Geriatric  (N = 5) | *d*  (95% CI) | Adj. *P-*Value | *d*  (95% CI) | Adj. *P-*Value | *d*  (95% CI) | Adj. *P-*Value |
| Male | 82.74  (26.67) | -21.59  (20.52) | 7.26  (5.90) | 4.06  (2.39, 5.73) | **<0.0001** | 3.17  (1.61, 4.73) | **<0.0001** | -1.63  (-2.95, -0.31) | **0.035** |

**Table S1. Group means and effect sizes for percent weight change across age groups.** Mean values with standard deviations (SD) are reported for percent weight change in adult, mid-age, and geriatric crickets overall and stratified by sex (N = number of individuals per group). Pairwise comparisons are quantified using Cohen’s *d* with Hedges’ *g* correction to account for small sample size bias. Effect sizes are reported alongside 95% confidence intervals (CI) in the format (lower, upper). Adults served as the reference group for adult-mid-age and adult-geriatric comparisons, while mid-age served as the reference group for mid-age-geriatric comparisons. Adjusted *P*-values were calculated using Tukey’s Honestly Significant Difference (HSD) post-hoc test.

| Percent Weight Change (%) | Mean (SD) | | *d*  (95% CI) | *P* |
| --- | --- | --- | --- | --- |
|  | Female | Male |  |  |
| Adult  (N_Female_ = 9, N_Male_ = 10) | 38.38  (20.74) | 82.74  (26.67) | -1.76  (-2.82, -0.70) | **<0.0001** |
| Mid-Age  (N_Female_ = 9, N_Male_ = 7) | 0.00  (16.35) | -21.59  (20.52) | 1.12  (0.06, 2.18) | 0.091 |
| Geriatric  (N_Female_ = 8, N_Male_ = 5) | -4.45  (9.97) | 7.26  (5.90) | -1.25  (-2.47, -0.03) | 0.87 |

**Table S2. Group means and effect sizes for percent weight change comparing sexes within each age group.** Mean values with standard deviations (SD) are reported for percent weight change in adult, mid-age, and geriatric crickets (N = number of individuals per group). Pairwise comparisons are quantified using Cohen’s *d* with Hedges’ *g* correction to account for small sample size bias. Effect sizes are reported alongside 95% confidence intervals (CI) in the format (lower, upper). Females served as the reference group for all comparisons. Adjusted *P*-values were calculated using Bonferroni’s multiple comparisons post-hoc test.

A. B.	C.

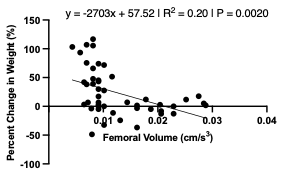

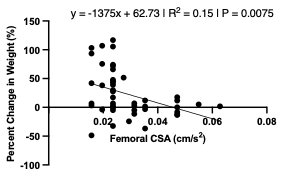

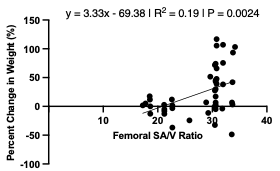

| **Femoral Traits** | **β** | **95% CI** | **R²** | ***P*** |
| --- | --- | --- | --- | --- |
| Volume (cm^3^) | -2703 | -4356 to -1050 | 0.20 | **0.0020** |
| CSA (cm/s^2^) | -1375 | -2363 to -386.7 | 0.15 | **0.0075** |
| SA/V | 3.33 | 1.24 to 5.41 | 0.19 | **0.0024** |

**Figure S1. Associations between femoral morphology and percent change in body weight. (A)** Femoral volume was negatively associated with percent weight change, with individuals possessing larger femoral volumes exhibiting greater weight loss. **(B)** Femoral cross-sectional area (CSA) similarly showed a negative relationship with weight change, indicating that crickets with thicker femora tended to lose more weight. **(C)** In contrast, the femoral surface area-to-volume (SA/V) ratio was positively associated with percent weight change, suggesting that individuals with relatively slender femora experienced less weight loss. Regression equations, coefficients (β), confidence intervals (95% CI), coefficient of determination (R²), and P-values are summarized in the accompanying table.

| **Source** | **SS** | **df** | **MS** | **F** | ***P*** |
| --- | --- | --- | --- | --- | --- |
| Regression | 55182 | 4 | 13795 | 30.45 | **<0.0001** |
| Group | 39918 | 2 | 19959 | 44.06 | **<0.0001** |
| Sex | 6585 | 1 | 6585 | 14.54 | **0.0005** |
| Femoral Volume | 3943 | 1 | 3943 | 8.71 | **0.0052** |
| Residual | 18573 | 41 | 453.0 |  |  |
| Total | 73755 | 45 |  |  |  |
| **Predictor** | **Estimate** | **Standard error** | | **95% CI** | ***P*** |
| Intercept | 25.43 | 11.13 | | 2.96 to 47.89 | **0.028** |
| Group[Mid-Age] | -89.97 | 10.99 | | -112.2 to -67.77 | **<0.0001** |
| Group[Geriatric] | -68.01 | 8.25 | | -84.67 to -51.35 | **<0.0001** |
| Sex[M] | 31.09 | 8.16 | | 14.62 to 47.56 | **0.0005** |
| Femoral Volume | 2435 | 825.4 | | 768.3 to 4102 | **0.0052** |

**Table S3. Effects of age, sex, and femoral volume on percent change in body weight.** ANCOVA revealed that age, sex, and femoral volume were significant predictors of weight change (R^2^ = 0.748). Using adult females as the reference group, mid-age and geriatric crickets lost more weight, males lost more than females, and greater femoral volume was associated with reduced weight loss. P-values were adjusted using the Holm-Bonferroni method.

| **Source** | **SS** | **df** | **MS** | **F** | ***P*** |
| --- | --- | --- | --- | --- | --- |
| Regression | 54113 | 4 | 13528 | 28.24 | **<0.0001** |
| Group | 41257 | 2 | 20629 | 43.06 | **<0.0001** |
| Sex | 5503 | 1 | 5503 | 11.49 | **0.0016** |
| Femoral CSA | 2875 | 1 | 2875 | 6.00 | **0.019** |
| Residual | 19642 | 41 | 479.1 |  |  |
| Total | 73755 | 45 |  |  |  |
| **Predictor** | **Estimate** | **Standard error** | | **95% CI** | ***P*** |
| Intercept | 24.14 | 13.34 | | -2.80 to 51.07 | **0.078** |
| Group[Mid-Age] | -83.49 | 10.44 | | -104.6 to -62.41 | **<0.0001** |
| Group[Geriatric] | -63.78 | 8.14 | | -80.21 to -47.35 | **<0.0001** |
| Sex[M] | 26.45 | 7.80 | | 10.69 to 42.20 | **0.0016** |
| Femoral CSA | 1080 | 441.0 | | 189.7 to 1971 | **0.019** |

**Table S4. Effects of age, sex, and femoral CSA on percent change in body weight.** ANCOVA showed that age group, sex, and femoral CSA were significant predictors of weight change (R² = 0.734). Using adult females as the reference group, mid-age and geriatric crickets lost more weight, males lost more than females, and larger femoral CSA was associated with reduced weight loss. P-values were adjusted using the Holm–Bonferroni method.

| **Source** | **SS** | **df** | **MS** | **F** | ***P*** |
| --- | --- | --- | --- | --- | --- |
| Regression | 54867 | 4 | 13717 | 29.77 | **<0.0001** |
| Group | 40142 | 2 | 20071 | 43.57 | **<0.0001** |
| Sex | 6333 | 1 | 6333 | 13.75 | **0.0006** |
| Femoral SA/V | 3628 | 1 | 3628 | 7.88 | **0.0076** |
| Residual | 18888 | 41 | 460.7 |  |  |
| Total | 73755 | 45 |  |  |  |
| **Predictor** | **Estimate** | **Standard error** | | **95% CI** | ***P*** |
| Intercept | 134.9 | 29.73 | | 74.87 to 194.9 | **<0.0001** |
| Group[Mid-Age] | -87.43 | 10.68 | | -109.0 to -65.85 | **<0.0001** |
| Group[Geriatric] | -67.43 | 8.29 | | -84.18 to -50.68 | **<0.0001** |
| Sex[M] | 30.52 | 8.23 | | 13.90 to 47.15 | **0.0006** |
| Femoral SA/V | -2.85 | 1.02 | | -4.90 to -0.80 | **0.0076** |

**Table S5. Effects of age, sex, and femoral SA/V ratio on percent change in body weight.** ANCOVA indicated that age group, sex, and femoral SA/V ratio were significant predictors of weight change (R² = 0.744). Using adult females as the reference group, mid-age and geriatric crickets lost more weight, males lost more than females, and higher femoral SA/V ratio was associated with greater weight loss. P-values were adjusted using the Holm–Bonferroni method.

**Appendix 5. Daily acquisition of associative learning / relationship to key morphological traits.**

| Overall | | | | | | | |
| --- | --- | --- | --- | --- | --- | --- | --- |
| Day | Group | N | Passed  (N, %) | Failed  (N, %) | Relative Risk (95% CI) | *P* | Adjusted *P*-value |
| 1 | Geriatric | 16 | 1 (6.25) | 15 (93.75) | Reference | − | − |
|  | Mid-Age | 19 | 7 (36.84) | 12 (63.16) | 5.89 (0.81, 42.99) | **0.047** | 0.094 |
|  | Adult | 20 | 9 (45.00) | 11 (55.00) | 7.20 (1.02, 51.05) | **0.022** | **0.044** |
| 2 | Geriatric | 14 | 5 (35.71) | 9 (64.29) | Reference | − | − |
|  | Mid-Age | 19 | 11 (57.89) | 8 (42.11) | 1.62 (0.73, 3.61) | 0.296 | 0.592 |
|  | Adult | 20 | 18 (90.00) | 2 (10.00) | 2.52 (1.23, 5.17) | **0.002** | **0.004** |
| 3 | Geriatric | 14 | 6 (42.86) | 8 (57.14) | Reference | − | − |
|  | Mid-Age | 18 | 15 (83.33) | 3 (16.67) | 1.94 (1.03, 3.68) | **0.027** | 0.054 |
|  | Adult | 20 | 20 (100) | 0 (0) | 2.28 (1.24, 4.18) | **0.0002** | **0.0004** |
| 4 | Geriatric | 14 | 7 (50.00) | 7 (50.00) | Reference | − | − |
|  | Mid-Age | 18 | 16 (88.89) | 2 (11.11) | 1.78 (1.03, 3.08) | **0.022** | **0.044** |
|  | Adult | 20 | 20 (100) | 0 (0) | 2.00 (1.15, 3.31) | **0.0006** | **0.0012** |
| 5 | Geriatric | 14 | 8 (57.14) | 6 (42.86) | Reference | - | - |
|  | Mid-Age | 18 | 18 (100) | 0 (0) | 1.70 (1.07, 1.70) | **0.0033** | **0.0066** |
|  | Adult | 20 | 20 (100) | 0 (0) | 1.71 (1.08, 2.70) | **0.0022** | **0.0044** |

| Female | | | | | | | | |
| --- | --- | --- | --- | --- | --- | --- | --- | --- |
| Day | Group | N | Passed  (N, %) | Failed  (N, %) | Relative Risk (95% CI) | | *P* | Adjusted *P*-value |
| 1 | Geriatric | 8 | 1 (12.50) | 7 (87.50) | Reference | | − | − |
|  | Mid-Age | 10 | 4 (40.00) | 6 (60.00) | 3.20 (0.44, 23.28) | | 0.314 | 0.628 |
|  | Adult | 10 | 5 (50.00) | 5 (50.00) | 4.00 (0.58, 27.71) | | 0.152 | 0.304 |
| 2 | Geriatric | 7 | 3 (42.86) | 4 (57.14) | Reference | | − | − |
|  | Mid-Age | 10 | 5 (50.00) | 5 (50.00) | 1.17 (0.41, 3.36) | | >0.99 | >0.99 |
|  | Adult | 10 | 9 (90.00) | 1 (10.00) | 2.10 (0.87, 5.06) | | 0.101 | 0.202 |
| 3 | Geriatric | 7 | 3 (42.86) | 4 (57.14) | Reference | | − | − |
|  | Mid-Age | 10 | 8 (80.00) | 2 (20.00) | 1.87 (0.75, 4.64) | | 0.162 | 0.324 |
|  | Adult | 10 | 10 (100) | 0 (0) | 2.22 (0.93, 5.28) | | **0.015** | **0.029** |
| 4 | Geriatric | 7 | 4 (57.14) | 3 (42.86) | Reference | | − | − |
|  | Mid-Age | 10 | 8 (80.00) | 2 (20.00) | 1.40 (0.69, 2.85) | | 0.593 | >0.99 |
|  | Adult | 10 | 10 (100) | 0 (0) | 1.67 (0.87, 3.21) | | 0.051 | 0.102 |
| 5 | Geriatric | 7 | 5 (71.43) | 2 (28.57) | Reference | | - | - |
|  | Mid-Age | 10 | 10 (100) | 0 (0) | 1.33 (0.82, 2.17) | | 0.154 | 0.308 |
|  | Adult | 10 | 10 (100) | 0 (0) | 1.33 (0.82, 2.17) | | 0.154 | 0.308 |
| Male | | | | | | | | |
| Day | Group | N | Passed  (N, %) | Failed  (N, %) | Relative Risk (95% CI) | *P* | | Adjusted *P*-value |
| 1 | Geriatric | 8 | 0 (0) | 8 (100) | Reference | − | | − |
|  | Mid-Age | 9 | 3 (33.33) | 6 (66.67) | 5.67 (0.33, 97.3) | 0.206 | | 0.412 |
|  | Adult | 10 | 4 (40.00) | 6 (60.00) | 6.80 (0.42, 111) | 0.092 | | 0.184 |
| 2 | Geriatric | 7 | 0 (0) | 7 (100) | Reference | − | | − |
|  | Mid-Age | 9 | 6 (66.67) | 3 (33.33) | 10.00 (0.66, 151) | **0.011** | | **0.022** |
|  | Adult | 10 | 9 (90.00) | 1 (10.00) | 13.50 (0.92, 198) | **0.0004** | | **0.0008** |
| 3 | Geriatric | 7 | 2 (28.57) | 5 (71.43) | Reference | − | | − |
|  | Mid-Age | 8 | 7 (87.50) | 1 (12.50) | 3.06 (0.92, 10.2) | **0.041** | | 0.082 |
|  | Adult | 10 | 10 (100) | 0 (0) | 3.33 (1.03, 10.8) | 0.0034 | | **0.0068** |
| 4 | Geriatric | 7 | 3 (42.86) | 4 (57.14) | Reference | − | | − |
|  | Mid-Age | 8 | 8 (100) | 0 (0) | 2.20 (0.92, 5.25) | **0.026** | | 0.052 |
|  | Adult | 10 | 10 (100) | 0 (0) | 2.22 (0.93, 5.28) | **0.015** | | **0.029** |
| 5 | Geriatric | 7 | 3 (42.86) | 4 (57.14) | Reference | - | | - |
|  | Mid-Age | 8 | 8 (100) | 0 (0) | 2.20 (0.92, 5.25) | **0.026** | | 0.052 |
|  | Adult | 10 | 10 (100) | 0 (0) | 2.22 (0.93, 5.28) | **0.015** | | **0.029** |

**Table S1. Daily escape paradigm success rates and effect size comparisons across age groups overall and stratified by sex.** For each experimental day, escape performance was reported for adult, mid-age, and geriatric crickets as the number and percentage of individuals that successfully completed the paradigm versus those that failed (N = number of individuals per group; % = percentage of group). Between-group comparisons were conducted using $\chi^{2}$ analysis or Fisher’s exact test, as appropriate. Effect sizes were expressed as relative risk (RR) with corresponding 95% confidence interval (CI) in the format (lower, upper). Geriatrics served as the reference group for all comparisons. Adjusted *P*-values were calculated using the Bonferroni correction.

| Day 1 | | | | |
| --- | --- | --- | --- | --- |
| Model | ^1^Mid-Age  OR [95% CI] | ^1^Mid-Age  *P*-value | ^1^Geriatric  OR [95% CI] | ^1^Geriatric  *P*-value |
| Group + Sex | 0.70 [0.19, 2.54] | 0.59 | 0.080 [0.00, 0.53] | **0.025** |
| + Antennal Length | 0.32 [0.03, 2.01] | 0.25 | 0.051 [0.00, 0.39] | **0.014** |
| + Antennal/Body Length Ratio | 0.41 [0.05, 2.74] | 0.37 | 0.057 [0.00, 0.46] | **0.020** |
| + Antennal/Weight Ratio | 0.85 [0.04, 15.9] | 0.91 | 0.10 [0.00, 1.74] | 0.14 |
| Day 2 | | | | |
| Model | ^1^Mid-Age  OR [95% CI] | ^1^Mid-Age  *P*-value | ^1^Geriatric  OR [95% CI] | ^1^Geriatric  *P*-value |
| Group + Sex | 0.15 [0.02, 0.75] | **0.033** | 0.062 [0.00, 0.33] | **0.0028** |
| + Antennal Length | 7.35 [1.07, 69.8] | 0.053 | 16.2 [2.89, 139] | **0.0036** |
| + Antennal/Body Length Ratio | 6.67 [0.89, 68.6] | 0.078 | 15.6 [2.59, 143] | **0.0057** |
| + Antennal/Weight Ratio | 8.94 [0.37, 377] | 0.20 | 20.0 [1.26, 618] | 0.052 |
| Day 3 | | | | |
| Model | ^1^Mid-Age  OR [95% CI] | ^1^Mid-Age  *P*-value | ^1^Geriatric  OR [95% CI] | ^1^Geriatric  *P*-value |
| Group + Sex | 0.12 [0.00, 1.33] | 0.089 | 0.02 [0.00, 0.19] | **0.00011** |
| + Antennal Length | 0.21 [0.00, 5.51] | 0.35 | 0.02 [0.00, 0.22] | **0.00028** |
| + Antennal/Body Length Ratio | 0.18 [0.00, 5.00] | 0.31 | 0.02 [0.00, 0.23] | **0.00044** |
| + Antennal/Weight Ratio | 0.05 [0.00, 5.74] | 0.24 | 0.01 [0.00, 0.30] | **0.0065** |
| Day 4 | | | | |
| Model | ^1^Mid-Age  OR [95% CI] | ^1^Mid-Age  *P*-value | ^1^Geriatric  OR [95% CI] | ^1^Geriatric  *P*-value |
| Group + Sex | 0. 17 [0.00, 2.23] | 0.19 | 0.02 [0.00, 0.22] | **0.00025** |
| + Antennal Length | 0.15 [0.00, 3.45] | 0.23 | 0.02 [0.00, 0.22] | **0.00035** |
| + Antennal/Body Length Ratio | 0.10 [0.00, 2.62] | 0.17 | 0.01 [0.00, 0.18] | **0.00028** |
| + Antennal/Weight Ratio | 0.01 [0.00, 2.04] | 0.097 | 0.00 [0.00, 0.16] | **0.0024** |
| Day 5 | | | | |
| Model | ^1^Mid-Age  OR [95% CI] | ^1^Mid-Age  *P*-value | ^1^Geriatric  OR [95% CI] | ^1^Geriatric  *P*-value |
| Group + Sex | 0.85 [0.00, 162] | 0.94 | 0.03 [0.00, 0.32] | **0.0014** |
| + Antennal Length | 0.32 [0.00, 61.9] | 0.59 | 0.02 [0.00, 0.28] | **0.00093** |
| + Antennal/Body Length Ratio | 0.24 [0.00, 50.8] | 0.52 | 0.02 [0.00, 0.23] | **0.00074** |
| + Antennal/Weight Ratio | 0.02 [0.00, 17.0] | 0.24 | 0.00 [0.00, 0.20] | **0.0040** |

**Table S2. Daily logistic regression models testing the effects of age, sex, and antennal investment on associative learning success.** Odds ratios (OR) with 95% confidence intervals (CI) are shown for mid-age and geriatric crickets relative to adult controls (reference = adult, female). Models were evaluated across five consecutive training days, beginning with age and sex, followed by inclusion of antennal length, antennal-to-body length ratio, and antennal-to-weight ratio as predictors. Values reflect the influence of age group and morphological traits on the likelihood of achieving task success on each day of training. P-values were adjusted using the Holm–Bonferroni method.

| **Overall** | Group [Mean (SD)] | | | Adult-Mid-Age | | Adult-Geriatric | | Mid-Age-Geriatric | |
| --- | --- | --- | --- | --- | --- | --- | --- | --- | --- |
|  | Adult  (N = 19-20) | Mid-Age  (N = 17-19) | Geriatric  (N = 16) | *d*  (95% CI) | Adj. *P-*Value | *d*  (95% CI) | Adj. *P-*Value | *d*  (95% CI) | Adj. *P-*Value |
| Main Trial |  |  |  |  |  |  |  |  |  |
| Decision Making Time (s) | 9.15  (2.75) | 20.54  (10.07) | 11.96  (4.45) | -1.53  (-2.24, -0.82) | **<0.0001** | -0.76  (-1.44, -0.08) | 0.32 | 1.05  (0.34, 1.75) | **0.0064** |
| Time to Punishment Arm (s) | 12.50  (6.15)  19 | 23.34  (12.33)  17 | 14.65  (6.63) | -1.11  (-1.91, -0.41) | **0.0016** | -0.33  (-1.00, 0.34) | 0.75 | 0.85  (0.14, 1.56) | **0.018** |
| Time to Reward Arm (s) | 8.54  (2.61) | 19.61  (10.64) | 10.93  (3.71) | -1.42  (-2.12, -0.71( | **<0.0001** | -0.74  (-1.42, -0.06) | 0.38 | 1.03  (0.32, 1.74) | **0.014** |
| Test Trial |  |  |  |  |  |  |  |  |  |
| Decision Making Time (s) | 13.73  (5.45) | 26.25  (16.89) | 16.11  (11.48) | -0.99  (-1.65, -0.32) | **0.0084** | -0.26  (-0.89, 0.37) | >0.99 | 0.69  (0.03, 1.34) | **0.012** |
| Time to Punishment Arm (s) | 13.26  (7.91) | 24.99 (27.87) | 12.39  (4.80) | -0.57  (-1.21, 0.07) | 0.50 | 0.13  (-0.53, 0.78) | >0.99 | 0.59  (-0.09, 1.27) | 0.53 |
| Time to Reward Arm (s) | 14.19  (6.22) | 27.52  (14.30) | 13.27  (5.92) | -1.20  (-1.88, -0.51) | **0.0063** | 0.15  (-0.51, 0.81) | >0.99 | 1.23  (0.51, 1.96) | **0.0010** |

**Table S3. Group means and effect sizes for decision-making measures across age groups.** Mean values with standard deviations (SD) are reported for each measure in adult, mid-age, and geriatric crickets (N = number of individuals per group). Pairwise comparisons are quantified using Cohen’s *d* with Hedges’ *g* correction to account for small sample size bias. Effect sizes are reported alongside 95% confidence intervals (CI) in the format (lower, upper). Adults served as the reference group for adult-mid-age and adult-geriatric comparisons, while mid-age served as the reference group for mid-age-geriatric comparisons. Adjusted *P*-values were calculated using Tukey’s Honestly Significant Difference (HSD) post-hoc test.

| **Female** | Group [Mean (SD)] | | | Adult-Mid-Age | | Adult-Geriatric | | Mid-Age-Geriatric | |
| --- | --- | --- | --- | --- | --- | --- | --- | --- | --- |
|  | Adult  (N = 9-10) | Mid-Age  (N = 9-10) | Geriatric  (N = 8) | *d*  (95% CI) | Adj. *P-*Value | *d*  (95% CI) | Adj. *P-*Value | *d*  (95% CI) | Adj. *P-*Value |
| Main Trial |  |  |  |  |  |  |  |  |  |
| Decision Making Time (s) | 9.05  (2.31) | 20.97  (10.29) | 12.64  (5.19) | -1.53  (-2.53, -0.53) | **0.0008** | -0.89  (-1.86, 0.08) | 0.51 | 0.94  (-0.04, 1.92) | **0.034** |
| Time to Punishment Arm (s) | 11.47  (5.15) | 22.38  (10.25) | 16.20  (7.65) | -1.28  (-2.30, -0.27) | **0.034** | -0.70  (-1.68, 0.28) | 0.53 | 0.64  (-0.33, 1.62) | 0.34 |
| Time to Reward Arm (s) | 8.48  (2.48) | 20.39  (11.54) | 11.28  (4.37) | -1.37  (-2.34, -0.39) | **0.0010** | -0.78  (-1.74, 0.19) | 0.67 | 0.95  (-0.03, 1.93) | **0.021** |
| Test Trial |  |  |  |  |  |  |  |  |  |
| Decision Making Time (s) | 12.09  (4.66) | 24.04  (17.18) | 14.45  (6.07) | -0.91  (-1.83, 0.01) | **0.046** | -0.42  (-1.36, 0.52) | 0.89 | 0.68  (-0.28, 1.63) | 0.16 |
| Time to Punishment Arm (s) | 11.22  (5.62) | 25.47  (33.84) | 13.59  (6.21) | -0.56  (-1.46, 0.33) | 0.18 | -0.38  (-1.32, 0.55) | 0.96 | 0.44  (-0.50, 1.38) | 0.34 |
| Time to Reward Arm (s) | 12.97  (7.05) | 22.61  (9.96) | 15.31  (7.27) | -1.07  (-2.01, -0.13) | 0.066 | -0.31  (-1.25, 0.62) | 0.86 | 0.78  (-0.18, 1.75) | 0.24 |
| **Male** | Group [Mean (SD)] | | | Adult-Mid-Age | | Adult-Geriatric | | Mid-Age-Geriatric | |
|  | Adult  (N = 10) | Mid-Age  (N = 8-9) | Geriatric  (N = 8) | *d*  (95% CI) | Adj. *P-*Value | *d*  (95% CI) | Adj. *P-*Value | *d*  (95% CI) | Adj. *P-*Value |
| Main Trial |  |  |  |  |  |  |  |  |  |
| Decision Making Time (s) | 9.25  (3.25) | 20.06  (10.43) | 11.28  (3.87) | -1.37  (-2.37, -0.37) | **0.0031** | -0.55  (-1.49, 0.40) | 0.80 | 1.03  (0.02, 2.05) | **0.028** |
| Time to Punishment Arm (s) | 13.43  (7.07) | 24.42  (14.99) | 13.10  (5.49) | -0.93  (-1.91, 0.05) | **0.034** | 0.05  (-1.11, 0.80) | >0.99 | 0.95  (-0.09, 1.98) | **0.039** |
| Time to Reward Arm (s) | 8.60  (2.86) | 18.73  (10.16) | 10.58  (3.18) | -1.33  (-2.33, -0.34) | **0.0071** | -0.63,  (-1.58, 0.32) | 0.82 | 1.00  (-0.01, 2.01) | **0.049** |
| Test Trial |  |  |  |  |  |  |  |  |  |
| Decision Making Time (s) | 15.36  (5.93) | 28.72  (17.23) | 11.21  (1.98) | -1.01  (-1.97, -0.06) | **0.028** | 0.85  (-0.12, 1.82) | 0.70 | 1.31  (0.26, 2.36) | **0.005** |
| Time to Punishment Arm (s) | 15.30  (9.56) | 24.45  (21.41) | 11.18  (2.74) | -0.54  (-1.45, 0.38) | 0.50 | 0.53  (-0.42, 1.48) | 0.88 | 0.80  (-0.19, 1.79) | 0.28 |
| Time to Reward Arm (s) | 15.42  (5.35) | 32.98  (16.87) | 11.23  (3.59) | -1.37  (-2.37, -0.37) | **0.0005** | 0.86  (-0.12, 1.83) | 0.62 | 1.64  (0.54, 2.74) | **<0.0001** |

**Table S4. Group means and effect sizes for decision-making measures across age groups stratified by sex.** Mean values with standard deviations (SD) are reported for each decision-making measure in juvenile, adult, and geriatric crickets (N = number of individuals per group). Pairwise comparisons are quantified using Cohen’s *d* with Hedges’ *g* correction to account for small sample size bias. Effect sizes are reported alongside 95% confidence intervals (CI) in the format (lower, upper). Adults served as the reference group for adult-mid-age and adult-geriatric comparisons, while mid-age served as the reference group for mid-age-geriatric comparisons. Adjusted *P*-values were calculated using Tukey’s Honestly Significant Difference (HSD) post-hoc test.

**Appendix 6. Average decision making / associations with select morphological traits.**

A.	B.	C.

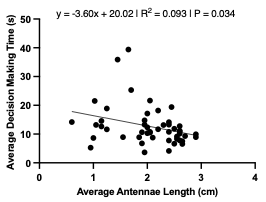

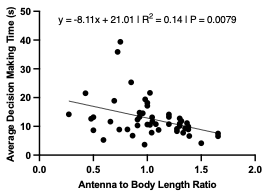

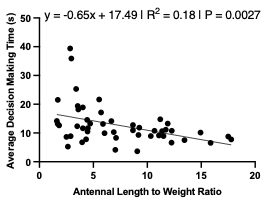

D.	E.	F.

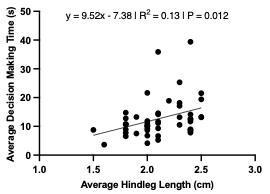

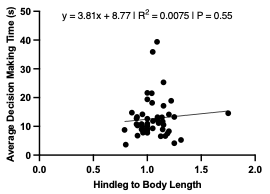

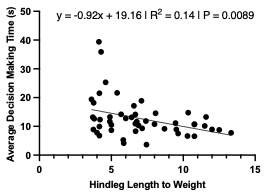

G.	H.	I.

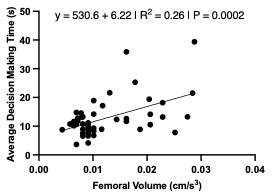

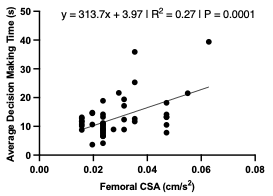

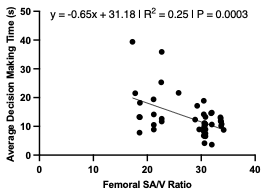

| **Trait** | **β** | **95% CI** | **R²** | ***P*** |
| --- | --- | --- | --- | --- |
| Antennal Length (cm) | -3.60 | -6.90 to -0.29 | 0.093 | **0.034** |
| Antennal-to-Body Length | -8.11 | -13.98 to -2.24 | 0.14 | **0.0079** |
| Antennal-to-Weight | -0.65 | -1.07 to -0.24 | 0.18 | **0.0027** |
| Hindleg Length (cm) | 9.52 | 2.17 to 16.87 | 0.13 | **0.012** |
| Hindleg-to-Body Length | 3.81 | -9.06 to 16.67 | 0.0075 | 0.55 |
| Hindleg-to-Weight | -0.92 | -1.60 to -0.24 | 0.14 | **0.0089** |
| Femoral Volume (cm^3^) | 530.6 | 263.8 to 797.4 | 0.26 | **0.0002** |
| Femoral CSA (cm/s^2^) | 313.7 | 161.1 to 466.3 | 0.27 | **0.0001** |
| Femoral SA/V | -0.65 | -0.99 to -0.32 | 0.25 | **0.0003** |

**Figure S1. Associations between morphological traits and decision time during the main escape learning trial.** Panels **(A–I)** depict simple linear regressions between decision time and individual morphological traits: **(A)** antennal length, **(B)** antennal-to-body length ratio, **(C)** antennal-to-weight ratio, **(D)** hindleg length, **(E)** hindleg-to-body length ratio, **(F)** hindleg-to-weight ratio, **(G)** femoral volume, **(H)** femoral cross-sectional area (CSA), and **(I)** femoral surface area-to-volume (SA/V) ratio. Regression coefficients (β), 95% confidence intervals, R² values, and P-values are summarized in the accompanying table. Traits with negative β values indicate shorter decision times with increasing trait size, whereas positive β values reflect longer decision times.

| **Source** | **SS** | **df** | **MS** | **F** | ***P*** |
| --- | --- | --- | --- | --- | --- |
| Regression | 824.0 | 4 | 206.0 | 6.26 | **0.0004** |
| Group | 423.1 | 2 | 211.6 | 6.43 | **0.0036** |
| Sex | 60.88 | 1 | 60.88 | 1.85 | 0.18 |
| Antennal Length | 25.08 | 1 | 25.08 | 0.76 | 0.39 |
| Residual | 1448 | 44 | 32.90 |  |  |
| Total | 2272 | 48 |  |  |  |
| **Predictor** | **Estimate** | **Standard error** | | **95% CI** | ***P*** |
| Intercept | 13.68 | 4.50 | | 4.61 to 22.75 | **0.0040** |
| Group[Mid-Age] | 8.25 | 2.35 | | 3.51 to 12.99 | **0.0011** |
| Group[Geriatric] | 2.37 | 2.04 | | -1.75 to 6.49 | 0.25 |
| Sex[M] | -2.37 | 1.74 | | -5.89 to 1.14 | 0.18 |
| Antennal Length | -1.47 | 1.68 | | -4.86 to 1.92 | 0.39 |

**Table S1. Effects of age, sex, and antennal length on main trial decision time during the escape learning task.** ANCOVA indicated that only age group significantly predicted decision time (R² = 0.363). Using adult females as the reference group, mid-age crickets exhibited longer decision times, while geriatric crickets did not differ. Neither sex nor antennal length was associated with variation in decision time. *P*-values were adjusted using the Holm–Bonferroni method.

| **Source** | **SS** | **df** | **MS** | **F** | ***P*** |
| --- | --- | --- | --- | --- | --- |
| Regression | 829.2 | 4 | 207.3 | 6.32 | **0.0004** |
| Group | 383.2 | 2 | 191.6 | 5.84 | **0.0056** |
| Sex | 52.80 | 1 | 52.80 | 1.61 | 0.21 |
| Antennal-to-Body Length | 30.19 | 1 | 30.19 | 0.92 | 0.34 |
| Residual | 1443 | 44 | 32.79 |  |  |
| Total | 2272 | 48 |  |  |  |
| **Predictor** | **Estimate** | **Standard error** | | **95% CI** | ***P*** |
| Intercept | 13.88 | 4.34 | | 5.13 to 22.62 | **0.0026** |
| Group[Mid-Age] | 7.96 | 2.45 | | 3.02 to 12.91 | **0.0023** |
| Group[Geriatric] | 2.04 | 2.15 | | -2.29 to 6.36 | 0.35 |
| Sex[M] | -2.12 | 1.67 | | -5.49 to 1.25 | 0.21 |
| Antennal-to-Body Length | -3.04 | 3.17 | | -9.43 to 3.35 | 0.34 |

**Table S2. Effects of age, sex, and antennal-to-body length ratio on main trial decision time during the escape learning task.** ANCOVA showed that age group was the only significant predictor of decision time (R² = 0.365). Using adult females as the reference group, mid-age crickets exhibited longer decision times, while geriatric crickets did not differ. Neither sex nor antennal-to-body length ratio was associated with decision time. *P*-values were adjusted using the Holm–Bonferroni method.

| **Source** | **SS** | **df** | **MS** | **F** | ***P*** |
| --- | --- | --- | --- | --- | --- |
| Regression | 802.8 | 4 | 200.7 | 6.01 | **0.0006** |
| Group | 383.2 | 2 | 191.6 | 5.74 | **0.0061** |
| Sex | 46.15 | 1 | 46.15 | 1.38 | 0.25 |
| Antennal-to-Weight | 3.81 | 1 | 3.81 | 0.11 | 0.74 |
| Residual | 1469 | 44 | 33.39 |  |  |
| Total | 2272 | 48 |  |  |  |
| **Predictor** | **Estimate** | **Standard error** | | **95% CI** | ***P*** |
| Intercept | 8.59 | 4.48 | | -0.44 to 17.61 | 0.062 |
| Group[Mid-Age] | 10.30 | 3.67 | | 2.90 to 17.71 | **0.0075** |
| Group[Geriatric] | 3.80 | 3.27 | | -2.79 to 10.38 | 0.25 |
| Sex[M] | -2.16 | 1.84 | | -5.87 to 1.54 | 0.25 |
| Antennal-to-Weight | 0.13 | 0.39 | | -0.65 to 0.91 | 0.74 |

**Table S3. Effects of age, sex, and antennal-to-weight ratio on main trial decision time during the escape learning task.** ANCOVA demonstrated that only age group significantly predicted decision time (R² = 0.353). Using adult females as the reference group, mid-age crickets exhibited longer decision times, whereas geriatric crickets did not differ. Neither sex nor antennal-to-weight ratio showed an association with decision time. *P*-values were adjusted using the Holm–Bonferroni method.

| **Source** | **SS** | **df** | **MS** | **F** | ***P*** |
| --- | --- | --- | --- | --- | --- |
| Regression | 825.0 | 4 | 206.3 | 6.27 | **0.0004** |
| Group | 534.8 | 2 | 267.4 | 8.13 | **0.0010** |
| Sex | 69.17 | 1 | 69.17 | 2.10 | 0.15 |
| Hindleg Length | 26.04 | 1 | 26.04 | 0.79 | 0.38 |
| Residual | 1447 | 44 | 32.88 |  |  |
| Total | 2272 | 48 |  |  |  |
| **Predictor** | **Estimate** | **Standard error** | | **95% CI** | ***P*** |
| Intercept | 19.41 | 10.69 | | -2.14 to 40.96 | 0.076 |
| Group[Mid-Age] | 10.95 | 2.78 | | 5.36 to 16.54 | **0.0003** |
| Group[Geriatric] | 3.86 | 2.22 | | -0.61 to 8.32 | 0.089 |
| Sex[M] | -2.95 | 2.03 | | -7.04 to 1.15 | 0.15 |
| Hindleg Length | -4.60 | 5.17 | | -15.02 to 5.82 | 0.38 |

**Table S4. Effects of age, sex, and hindleg length on main trial decision time during the escape learning task.** ANCOVA indicated that age group significantly predicted decision time (R² = 0.363). Using adult females as the reference group, mid-age crickets exhibited longer decision times, while geriatric crickets did not differ. Neither sex nor hindleg length was associated with variation in decision time. *P*-values were adjusted using the Holm–Bonferroni method.

| **Source** | **SS** | **df** | **MS** | **F** | ***P*** |
| --- | --- | --- | --- | --- | --- |
| Regression | 845.3 | 4 | 211.3 | 6.52 | **0.0003** |
| Group | 530.7 | 2 | 265.3 | 8.18 | **0.0010** |
| Sex | 89.63 | 1 | 89.63 | 2.76 | 0.10 |
| Hindleg-to-Weight | 46.37 | 1 | 46.37 | 1.43 | 0.24 |
| Residual | 1426 | 44 | 32.42 |  |  |
| Total | 2272 | 48 |  |  |  |
| **Predictor** | **Estimate** | **Standard error** | | **95% CI** | ***P*** |
| Intercept | 2.59 | 6.39 | | -10.30 to 15.47 | 0.69 |
| Group[Mid-Age] | 13.04 | 3.74 | | 5.50 to 20.58 | **0.0011** |
| Group[Geriatric] | 6.63 | 3.66 | | -0.75 to 14.01 | 0.077 |
| Sex[M] | -3.88 | 2.34 | | -8.59 to 0.82 | 0.10 |
| Hindleg-to-Weight | 0.86 | 0.72 | | -0.59 to 2.32 | 0.24 |

**Table S5. Effects of age, sex, and hindleg-to-weight ratio on main trial decision time during the escape learning task.** ANCOVA revealed that age group was the only significant predictor of decision time (R² = 0.372). Using adult females as the reference group, mid-age crickets exhibited longer decision times, while geriatric crickets did not differ. Neither sex nor hindleg-to-weight ratio was associated with decision time. *P*-values were adjusted using the Holm–Bonferroni method.

| **Source** | **SS** | **df** | **MS** | **F** | ***P*** |
| --- | --- | --- | --- | --- | --- |
| Regression | 803.6 | 4 | 200.9 | 6.12 | **0.0005** |
| Group | 209.6 | 2 | 104.8 | 3.19 | 0.051 |
| Sex | 1.44 | 1 | 1.44 | 0.04 | 0.84 |
| Femoral Volume | 25.22 | 1 | 25.22 | 0.77 | 0.39 |
| Residual | 1411 | 43 | 32.82 |  |  |
| Total | 2215 | 47 |  |  |  |
| **Predictor** | **Estimate** | **Standard error** | | **95% CI** | ***P*** |
| Intercept | 7.65 | 2.98 | | 1.63 to 13.66 | **0.014** |
| Group[Mid-Age] | 7.45 | 2.95 | | 1.49 to 13.40 | **0.016** |
| Group[Geriatric] | 2.63 | 2.13 | | -1.67 to 6.92 | 0.22 |
| Sex[M] | -0.45 | 2.16 | | -4.80 to 3.90 | 0.84 |
| Femoral Volume | 194.1 | 221.5 | | -252.5 to 640.7 | 0.39 |

**Table S6. Effects of age, sex, and femoral volume on decision time during the escape learning task.** ANCOVA showed that only age group approached significance as a predictor of decision time (R² = 0.363). Using adult females as the reference group, mid-age crickets exhibited longer decision times, whereas geriatric crickets did not differ. Neither sex nor femoral volume influenced decision time. *P*-values were adjusted using the Holm–Bonferroni method.

| **Source** | **SS** | **df** | **MS** | **F** | ***P*** |
| --- | --- | --- | --- | --- | --- |
| Regression | 832.4 | 4 | 208.1 | 6.47 | **0.0004** |
| Group | 218.9 | 2 | 109.5 | 3.40 | **0.042** |
| Sex | 0.40 | 1 | 0.40 | 0.013 | 0.91 |
| Femoral CSA | 54.01 | 1 | 54.01 | 1.68 | 0.20 |
| Residual | 1383 | 43 | 32.16 |  |  |
| Total | 2215 | 47 |  |  |  |
| **Predictor** | **Estimate** | **Standard error** | | **95% CI** | ***P*** |
| Intercept | 5.89 | 3.44 | | -1.04 to 12.83 | 0.094 |
| Group[Mid-Age] | 7.01 | 2.69 | | 1.58 to 12.44 | **0.013** |
| Group[Geriatric] | 2.75 | 2.01 | | -1.30 to 6.80 | 0.18 |
| Sex[M] | -0.22 | 1.99 | | -4.23 to 3.78 | 0.91 |
| Femoral CSA | 147.0 | 113.4 | | -81.74 to 375.7 | 0.20 |

**Table S7. Effects of age, sex, and femoral CSA on decision time during the escape learning task.** ANCOVA indicated that age group significantly predicted decision time (R² = 0.376). Using adult females as the reference group, mid-age crickets exhibited longer decision times, while geriatric crickets did not differ. Neither sex nor femoral CSA was associated with decision time. *P*-values were adjusted using the Holm–Bonferroni method.

| **Source** | **SS** | **df** | **MS** | **F** | ***P*** |
| --- | --- | --- | --- | --- | --- |
| Regression | 809.2 | 4 | 202.3 | 6.19 | **0.0005** |
| Group | 224.7 | 2 | 112.3 | 3.44 | **0.041** |
| Sex | 0.64 | 1 | 0.64 | 0.02 | 0.89 |
| Femoral SA/V | 30.87 | 1 | 30.87 | 0.94 | 0.34 |
| Residual | 1406 | 43 | 32.69 |  |  |
| Total | 2215 | 47 |  |  |  |
| **Predictor** | **Estimate** | **Standard error** | | **95% CI** | ***P*** |
| Intercept | 17.32 | 7.84 | | 1.51 to 33.14 | **0.033** |
| Group[Mid-Age] | 7.41 | 2.83 | | 1.70 to 13.12 | **0.012** |
| Group[Geriatric] | 2.62 | 2.10 | | -1.63 to 6.86 | 0.22 |
| Sex[M] | -0.30 | 2.17 | | -4.67 to 4.07 | 0.89 |
| Femoral SA/V | -0.26 | 0.27 | | -0.80 to 0.28 | 0.34 |

**Table S8. Effects of age, sex, and femoral SA/V ratio on decision time during the escape learning task.** ANCOVA revealed that age group significantly predicted decision time (R² = 0.365). Using adult females as the reference group, mid-age crickets exhibited longer decision times, while geriatric crickets did not differ. Neither sex nor femoral SA/V ratio was associated with variation in decision time. *P*-values were adjusted using the Holm–Bonferroni method.

A.	B.	C.

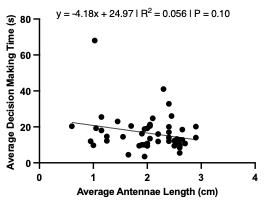

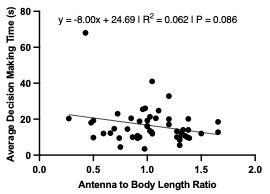

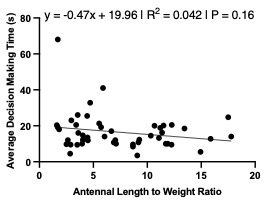

D.	E.	F.

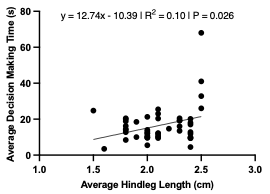

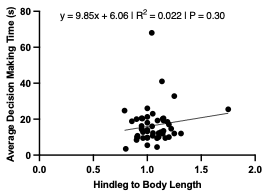

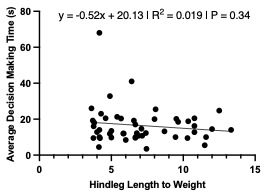

G.	H.	I.

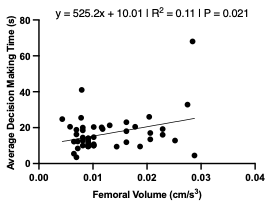

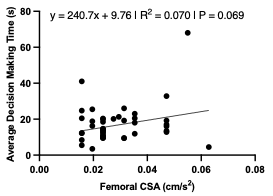

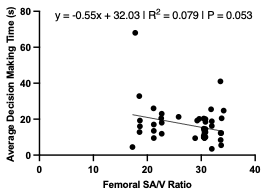

| **Trait** | **β** | **95% CI** | **R²** | ***P*** |
| --- | --- | --- | --- | --- |
| Antennal Length (cm) | -4.18 | -9.21 to 0.86 | 0.056 | 0.10 |
| Antennal-to-Body Length | -8.00 | -17.17 to 1.17 | 0.062 | 0.086 |
| Antennal-to-Weight | -0.47 | -1.14 to 0.19 | 0.042 | 0.16 |
| Hindleg Length (cm) | 12.74 | 1.61 to 23.87 | 0.10 | **0.026** |
| Hindleg-to-Body Length | 9.85 | -9.21 to 28.91 | 0.022 | 0.30 |
| Hindleg-to-Weight | -0.52 | -1.60 to 0.56 | 0.019 | 0.34 |
| Femoral Volume (cm^3^) | 525.2 | 84.42 to 965.9 | 0.11 | **0.021** |
| Femoral CSA (cm/s^2^) | 240.7 | -19.39 to 500.8 | 0.070 | 0.069 |
| Femoral SA/V | -0.55 | -1.11 to 0.01 | 0.079 | 0.053 |

**Figure S2. Relationships between morphological characteristics and decision time during the test escape learning trial.** Panels **(A–I)** illustrate simple linear regressions assessing the link between decision time and each morphological parameter: **(A)** antennal length, **(B)** antennal-to-body length ratio, **(C)** antennal-to-weight ratio, **(D)** hindleg length, **(E)** hindleg-to-body length ratio, **(F)** hindleg-to-weight ratio, **(G)** femoral volume, **(H)** femoral CSA, and **(I)** femoral SA/V ratio. Corresponding regression slopes (β), 95% confidence intervals, R² values, and P-values are provided in the table. Negative β values denote faster decision times with increasing trait size, whereas positive β values indicate the opposite pattern.

| **Source** | **SS** | **df** | **MS** | **F** | ***P*** |
| --- | --- | --- | --- | --- | --- |
| Regression | 1059 | 4 | 264.7 | 0.90 | 0.47 |
| Group | 392.7 | 2 | 196.3 | 0.67 | 0.52 |
| Sex | 4.48 | 1 | 4.48 | 0.02 | 0.90 |
| Hindleg Length | 178.1 | 1 | 178.1 | 0.61 | 0.44 |
| Residual | 12882 | 44 | 292.8 |  |  |
| Total | 13941 | 48 |  |  |  |
| **Predictor** | **Estimate** | **Standard error** | | **95% CI** | ***P*** |
| Intercept | -9.96 | 31.91 | | -74.26 to 54.34 | 0.76 |
| Group[Mid-Age] | 3.79 | 8.28 | | -12.90 to 20.47 | 0.65 |
| Group[Geriatric] | -3.75 | 6.61 | | -17.08 to 9.58 | 0.57 |
| Sex[M] | 0.75 | 6.06 | | -11.46 to 12.96 | 0.90 |
| Hindleg Length | 12.03 | 15.43 | | -19.06 to 43.12 | 0.44 |

**Table S9. Effects of age, sex, and hindleg length on decision time during the test escape learning trial.** ANCOVA showed that neither age group, sex, nor hindleg length significantly predicted decision time (R² = 0.08). Using adult females as the reference group, neither mid-age nor geriatric crickets differed in performance, and hindleg length was not associated with variation in decision time. *P*-values were adjusted using the Holm–Bonferroni method.

| **Source** | **SS** | **df** | **MS** | **F** | ***P*** |
| --- | --- | --- | --- | --- | --- |
| Regression | 1187 | 4 | 296.8 | 3.31 | **0.019** |
| Group | 479.6 | 2 | 239.8 | 2.67 | 0.081 |
| Sex | 27.68 | 1 | 27.68 | 0.31 | 0.58 |
| Femoral Volume | 47.19 | 1 | 47.19 | 0.53 | 0.47 |
| Residual | 3857 | 43 | 89.70 |  |  |
| Total | 5044 | 47 |  |  |  |
| **Predictor** | **Estimate** | **Standard error** | | **95% CI** | ***P*** |
| Intercept | 10.72 | 4.93 | | 0.78 to 20.66 | **0.035** |
| Group[Mid-Age] | 7.71 | 4.88 | | -2.14 to 17.55 | 0.12 |
| Group[Geriatric] | -1.97 | 3.52 | | -9.07 to 5.14 | 0.58 |
| Sex[M] | 1.98 | 3.56 | | -5.21 to 9.17 | 0.58 |
| Femoral Volume | 265.5 | 366.1 | | -472.7 to 1004 | 0.47 |

**Table S10. Effects of age, sex, and femoral volume on decision time during the test escape learning trial.** ANCOVA indicated that the overall model explained 23.5% of the variance in decision time (R² = 0.235), but none of the predictors reached statistical significance after adjustment. Using adult females as the reference group, mid-age and geriatric crickets did not differ in decision time, and neither sex nor femoral volume showed an association with performance. *P*-values were adjusted using the Holm–Bonferroni method.

A.	B.	C.

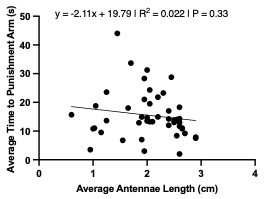

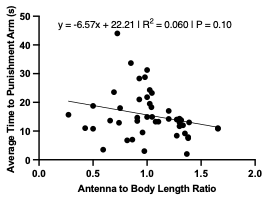

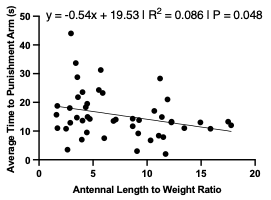

D.	E.	F.

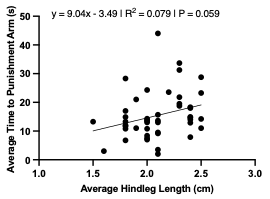

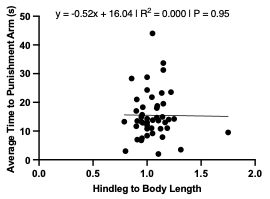

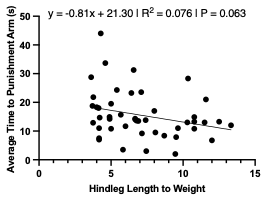

G.	H.	I.

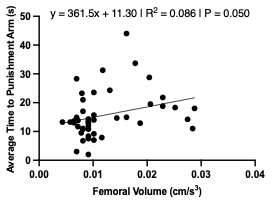

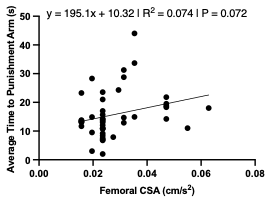

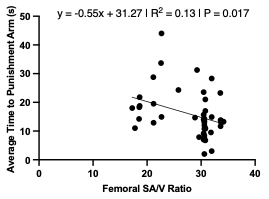

| **Trait** | **β** | **95% CI** | **R²** | ***P*** |
| --- | --- | --- | --- | --- |
| Antennal Length (cm) | -2.11 | -6.39 to 2.17 | 0.022 | 0.33 |
| Antennal-to-Body Length | -6.57 | -14.44 to 1.30 | 0.060 | 0.10 |
| Antennal-to-Weight | -0.54 | -1.08 to -0.00 | 0.086 | **0.048** |
| Hindleg Length (cm) | 9.04 | -0.34 to 18.42 | 0.079 | 0.059 |
| Hindleg-to-Body Length | -0.52 | -16.52 to 15.48 | 0.000 | 0.95 |
| Hindleg-to-Weight | -0.81 | -1.67 to 0.05 | 0.076 | 0.063 |
| Femoral Volume (cm^3^) | 361.5 | -0.25 to 723.2 | 0.086 | 0.050 |
| Femoral CSA (cm/s^2^) | 195.1 | -17.92 to 408.2 | 0.074 | 0.072 |
| Femoral SA/V | -0.55 | -1.00 to -0.10 | 0.13 | **0.017** |

**Figure S3. Associations between morphological traits and time spent traveling to the punishment arm during main escape learning trials.** Panels **(A–I)** illustrate simple linear regressions relating punishment arm time to individual morphological parameter: **(A)** antennal length, **(B)** antennal-to-body length ratio, **(C)** antennal-to-weight ratio, **(D)** hindleg length, **(E)** hindleg-to-body length ratio, **(F)** hindleg-to-weight ratio, **(G)** femoral volume, **(H)** femoral CSA, and **(I)** femoral SA/V ratio. Regression coefficients (β), 95% confidence intervals, R² values, and *P*-values are presented in the table. Negative β values indicate reduced punishment arm time with increasing trait size, while positive β values indicate longer time in the punishment arm.

| **Source** | **SS** | **df** | **MS** | **F** | ***P*** |
| --- | --- | --- | --- | --- | --- |
| Regression | 683.8 | 4 | 170.9 | 2.92 | **0.033** |
| Group | 415.2 | 2 | 207.6 | 3.55 | **0.038** |
| Sex | 10.53 | 1 | 10.53 | 0.18 | 0.67 |
| Antennal-to-Weight | 15.90 | 1 | 15.90 | 0.27 | 0.61 |
| Residual | 2401 | 41 | 58.57 |  |  |
| Total | 3085 | 45 |  |  |  |
| **Predictor** | **Estimate** | **Standard error** | | **95% CI** | ***P*** |
| Intercept | 9.332 | 6.42 | | -3.64 to 22.31 | 0.15 |
| Group[Mid-Age] | 11.65 | 5.28 | | 0.99 to 22.31 | **0.033** |
| Group[Geriatric] | 4.42 | 4.59 | | -4.85 to 13.69 | 0.34 |
| Sex[M] | -1.04 | 2.46 | | -6.02 to 3.93 | 0.67 |
| Antennal-to-Weight | 0.28 | 0.54 | | -0.80 to 1.36 | 0.61 |

**Table S11. Effects of age, sex, and antennal-to-weight ratio on punishment arm time during the main escape learning trial.** ANCOVA revealed that age group significantly predicted punishment arm time (R² = 0.222). Using adult females as the reference group, mid-age crickets spent more time in the punishment arm, whereas geriatric crickets did not differ. Neither sex nor antennal-to-weight ratio showed an association with punishment arm time. *P*-values were adjusted using the Holm–Bonferroni method.

| **Source** | **SS** | **df** | **MS** | **F** | ***P*** |
| --- | --- | --- | --- | --- | --- |
| Regression | 684.0 | 4 | 171.0 | 3.03 | **0.028** |
| Group | 223.5 | 2 | 111.8 | 1.98 | 0.15 |
| Sex | 14.49 | 1 | 14.49 | 0.26 | 0.61 |
| Femoral SA/V | 34.26 | 1 | 34.26 | 0.61 | 0.44 |
| Residual | 2254 | 40 | 56.35 |  |  |
| Total | 2938 | 44 |  |  |  |
| **Predictor** | **Estimate** | **Standard error** | | **95% CI** | ***P*** |
| Intercept | 20.27 | 10.61 | | -1.16 to 41.71 | 0.063 |
| Group[Mid-Age] | 7.52 | 3.79 | | -0.14 to 15.17 | 0.054 |
| Group[Geriatric] | 2.52 | 2.79 | | -3.12 to 8.15 | 0.37 |
| Sex[M] | 1.56 | 3.07 | | -4.65 to 7.76 | 0.61 |
| Femoral SA/V | -0.29 | 0.37 | | -1.03 to 0.46 | 0.44 |

**Table S12. Effects of age, sex, and femoral SA/V ratio on punishment arm time during the main escape learning trial.** ANCOVA showed that the model explained 23.3% of the variance in punishment arm time (R² = 0.233). Neither age group, sex, nor femoral SA/V ratio significantly predicted punishment arm time after adjustment. Using adult females as the reference group, mid-age crickets showed a nonsignificant trend toward longer punishment arm times. *P*-values were adjusted using the Holm–Bonferroni method.

A.	B.	C.

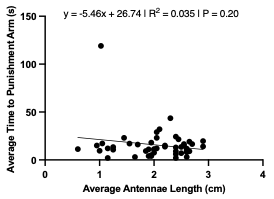

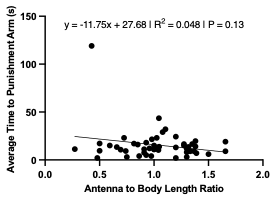

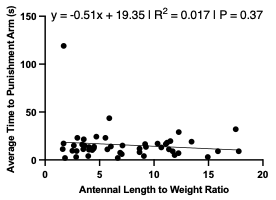

D.	E.	F.

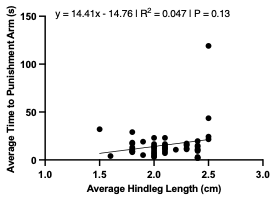

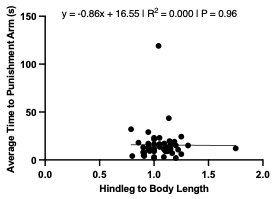

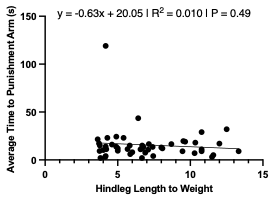

G.	H.	I.

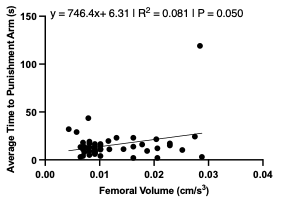

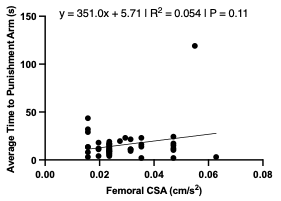

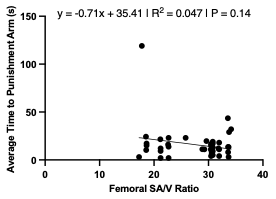

| **Trait** | **β** | **95% CI** | **R²** | ***P*** |
| --- | --- | --- | --- | --- |
| Antennal Length (cm) | -5.46 | -13.91 to 2.98 | 0.035 | 0.20 |
| Antennal-to-Body Length | -11.75 | -27.06 to 3.57 | 0.048 | 0.13 |
| Antennal-to-Weight | -0.51 | -1.63 to 0.61 | 0.017 | 0.37 |
| Hindleg Length (cm) | 14.41 | -4.60 to 33.42 | 0.047 | 0.13 |
| Hindleg-to-Body Length | -0.86 | -32.84 to 31.12 | 0.000 | 0.96 |
| Hindleg-to-Weight | -0.63 | -2.43 to 1.17 | 0.010 | 0.49 |
| Femoral Volume (cm^3^) | 746.4 | 1.46 to 1491 | 0.081 | 0.050 |
| Femoral CSA (cm/s^2^) | 351.0 | -85.17 to 787.2 | 0.054 | 0.11 |
| Femoral SA/V | -0.71 | -1.65 to 0.24 | 0.047 | 0.14 |

**Figure S4. Associations between morphological traits and time spent traveling to the punishment arm during the test escape learning trial.** Panels **(A–I)** display simple linear regressions between punishment arm time and individual morphological measures: **(A)** antennal length, **(B)** antennal-to-body length ratio, **(C)** antennal-to-weight ratio, **(D)** hindleg length, **(E)** hindleg-to-body length ratio, **(F)** hindleg-to-weight ratio, **(G)** femoral volume, **(H)** femoral CSA, and **(I)** femoral SA/V ratio. Regression coefficients (β), 95% confidence intervals, R² values, and *P*-values are summarized in the accompanying table. Traits with negative β values indicate shorter punishment arm times with increasing trait size, whereas positive β values reflect the opposite pattern.

A.	B.	C.

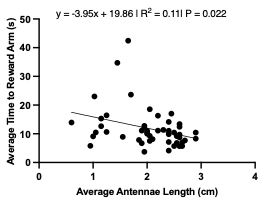

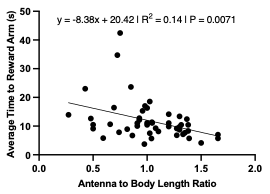

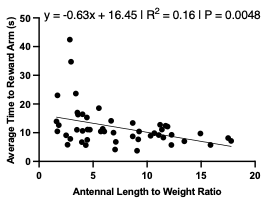

D.	E.	F.

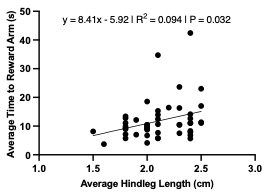

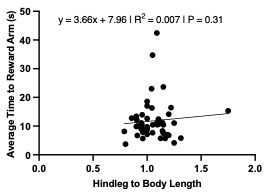

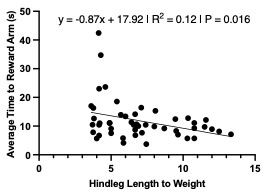

G.	H.	I.

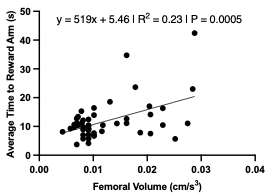

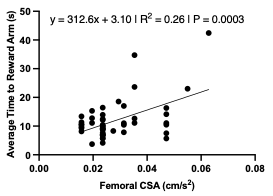

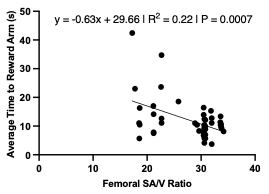

| **Trait** | **β** | **95% CI** | **R²** | ***P*** |
| --- | --- | --- | --- | --- |
| Antennal Length (cm) | -3.95 | -7.30 to -0.60 | 0.11 | **0.022** |
| Antennal-to-Body Length | -8.38 | -14.37 to -2.39 | 0.14 | **0.0071** |
| Antennal-to-Weight | -0.63 | -1.06 to -0.20 | 0.16 | **0.0048** |
| Hindleg Length (cm) | 8.41 | 0.77 to 16.06 | 0.094 | **0.032** |
| Hindleg-to-Body Length | 3.66 | -9.49 to 16.82 | 0.007 | 0.31 |
| Hindleg-to-Weight | -0.87 | -1.57 to -0.17 | 0.12 | **0.016** |
| Femoral Volume (cm^3^) | 519 | 240.6 to 797.5 | 0.23 | **0.0005** |
| Femoral CSA (cm/s^2^) | 312.6 | 154.1 to 471.0 | 0.26 | **0.0003** |
| Femoral SA/V | -0.63 | -0.98 to -0.28 | 0.22 | **0.0007** |

**Figure S5. Linear relationships between morphological traits and time spent traveling to the reward arm during the main escape learning trial.** Panels **(A–I)** display simple linear regressions between reward arm time and individual morphological measures: **(A)** antennal length, **(B)** antennal-to-body length ratio, **(C)** antennal-to-weight ratio, **(D)** hindleg length, **(E)** hindleg-to-body length ratio, **(F)** hindleg-to-weight ratio, **(G)** femoral volume, **(H)** femoral CSA, and **(I)** femoral SA/V ratio. Regression equations, slope (β), 95% confidence intervals, R² values, and *P*-values are summarized in the accompanying table. Negative slopes indicate decreased reward arm time with increasing trait value, while positive slopes reflect the opposite trend.

| **Source** | **SS** | **df** | **MS** | **F** | ***P*** |
| --- | --- | --- | --- | --- | --- |
| Regression | 789.5 | 4 | 197.4 | 5.48 | **0.0011** |
| Group | 336.7 | 2 | 168.3 | 4.68 | **0.014** |
| Sex | 76.56 | 1 | 76.56 | 2.13 | 0.15 |
| Antennal Length | 57.28 | 1 | 57.28 | 1.59 | 0.21 |
| Residual | 1585 | 44 | 36.01 |  |  |
| Total | 2374 | 48 |  |  |  |
| **Predictor** | **Estimate** | **Standard error** | | **95% CI** | ***P*** |
| Intercept | 15.00 | 4.71 | | 5.51 to 24.48 | **0.0027** |
| Group[Mid-Age] | 7.22 | 2.46 | | 2.26 to 12.18 | **0.0053** |
| Group[Geriatric] | 1.64 | 2.14 | | -2.67 to 5.95 | 0.45 |
| Sex[M] | -2.66 | 1.83 | | -6.34 to 1.02 | 0.15 |
| Antennal Length | -2.22 | 1.76 | | -5.77 to 1.33 | 0.21 |

**Table S13. Effects of age, sex, and antennal length on time spent in the reward arm during the main escape learning trial.** ANCOVA demonstrated that age group significantly predicted reward arm time (R² = 0.333). Using adult females as the reference group, mid-age crickets spent more time traveling to the reward arm, while geriatric crickets did not differ. Neither sex nor antennal length was associated with variation in reward arm time. *P*-values were adjusted using the Holm–Bonferroni method.

| **Source** | **SS** | **df** | **MS** | **F** | ***P*** |
| --- | --- | --- | --- | --- | --- |
| Regression | 786.1 | 4 | 196.5 | 5.45 | **0.0012** |
| Group | 317.4 | 2 | 158.7 | 4.40 | **0.018** |
| Sex | 59.02 | 1 | 59.02 | 1.64 | 0.21 |
| Antennal-to-Body Length | 53.91 | 1 | 53.91 | 1.49 | 0.23 |
| Residual | 1588 | 44 | 36.09 |  |  |
| Total | 2374 | 48 |  |  |  |
| **Predictor** | **Estimate** | **Standard error** | | **95% CI** | ***P*** |
| Intercept | 14.62 | 4.55 | | 5.45 to 23.80 | **0.0025** |
| Group[Mid-Age] | 7.01 | 2.58 | | 1.82 to 12.20 | **0.0092** |
| Group[Geriatric] | 1.30 | 2.25 | | -3.24 to 5.83 | 0.57 |
| Sex[M] | -2.24 | 1.75 | | -5.78 to 1.29 | 0.21 |
| Antennal-to-Body Length | -4.06 | 3.33 | | -10.76 to 2.64 | 0.23 |

**Table S14. Effects of age, sex, and antennal-to-body length ratio on travel time to the reward arm during the main escape learning trial.** ANCOVA revealed that age group significantly predicted reward arm travel time (R² = 0.331). Using adult females as the reference group, mid-age crickets required more time to reach the reward arm, while geriatric crickets did not differ. Neither sex nor antennal-to-body length ratio was associated with travel time. *P*-values were adjusted using the Holm–Bonferroni method.

| **Source** | **SS** | **df** | **MS** | **F** | ***P*** |
| --- | --- | --- | --- | --- | --- |
| Regression | 732.7 | 4 | 183.2 | 4.91 | **0.0023** |
| Group | 338.2 | 2 | 169.1 | 4.53 | **0.016** |
| Sex | 41.04 | 1 | 41.04 | 1.10 | 0.30 |
| Antennal-to-Weight | 0.44 | 1 | 0.44 | 0.012 | 0.91 |
| Residual | 1641 | 44 | 37.31 |  |  |
| Total | 2374 | 48 |  |  |  |
| **Predictor** | **Estimate** | **Standard error** | | **95% CI** | ***P*** |
| Intercept | 8.96 | 4.73 | | -0.58 to 18.50 | 0.065 |
| Group[Mid-Age] | 9.12 | 3.88 | | 1.29 to 16.94 | **0.023** |
| Group[Geriatric] | 2.76 | 3.46 | | -4.20 to 9.73 | 0.43 |
| Sex[M] | -2.04 | 1.94 | | -5.96 to 1.88 | 0.30 |
| Antennal-to-Weight | 0.04 | 0.41 | | -0.78 to 0.87 | 0.91 |

**Table S15. Effects of age, sex, and antennal-to-weight ratio on travel time to the reward arm during the main escape learning trial.** ANCOVA showed that age group significantly predicted reward arm travel time (R² = 0.353). Using adult females as the reference group, mid-age crickets took longer to reach the reward arm, whereas geriatric crickets did not differ. Neither sex nor antennal-to-weight ratio was associated with travel time. *P*-values were adjusted using the Holm–Bonferroni method.

| **Source** | **SS** | **df** | **MS** | **F** | ***P*** |
| --- | --- | --- | --- | --- | --- |
| Regression | 778.7 | 4 | 194.7 | 5.37 | **0.0013** |
| Group | 545.2 | 2 | 272.6 | 7.52 | **0.0016** |
| Sex | 89.12 | 1 | 89.12 | 2.46 | 0.12 |
| Hindleg Length | 46.48 | 1 | 46.48 | 1.28 | 0.26 |
| Residual | 1595 | 44 | 36.26 |  |  |
| Total | 2374 | 48 |  |  |  |
| **Predictor** | **Estimate** | **Standard error** | | **95% CI** | ***P*** |
| Intercept | 22.01 | 11.23 | | -0.62 to 44.64 | 0.056 |
| Group[Mid-Age] | 11.01 | 2.91 | | 5.13 to 16.88 | **0.0005** |
| Group[Geriatric] | 3.72 | 2.33 | | -0.97 to 8.42 | 0.12 |
| Sex[M] | -3.34 | 2.13 | | -7.64 to 0.95 | 0.12 |
| Hindleg Length | -6.15 | 5.43 | | -17.09 to 4.80 | 0.26 |

**Table S16. Effects of age, sex, and hindleg length on travel time to the reward arm during the main escape learning trial.** ANCOVA indicated that age group significantly predicted reward arm travel time (R² = 0.328). Using adult females as the reference group, mid-age crickets took longer to reach the reward arm, while geriatric crickets did not differ. Neither sex nor hindleg length was associated with travel time. *P*-values were adjusted using the Holm–Bonferroni method.

| **Source** | **SS** | **df** | **MS** | **F** | ***P*** |
| --- | --- | --- | --- | --- | --- |
| Regression | 770.8 | 4 | 192.7 | 5.29 | **0.0014** |
| Group | 487.5 | 2 | 243.7 | 6.69 | **0.0029** |
| Sex | 83.83 | 1 | 83.83 | 2.30 | 0.14 |
| Hindleg-to-Weight | 38.52 | 1 | 38.52 | 1.06 | 0.31 |
| Residual | 1603 | 44 | 36.44 |  |  |
| Total | 2374 | 48 |  |  |  |
| **Predictor** | **Estimate** | **Standard error** | | **95% CI** | ***P*** |
| Intercept | 2.68 | 6.78 | | -10.98 to 16.34 | 0.69 |
| Group[Mid-Age] | 12.20 | 3.97 | | 4.20 to 20.19 | **0.0036** |
| Group[Geriatric] | 5.85 | 3.88 | | -1.97 to 13.68 | 0.14 |
| Sex[M] | -3.76 | 2.48 | | -8.75 to 1.24 | 0.14 |
| Hindleg-to-Weight | 0.79 | 0.76 | | -0.75 to 2.33 | 0.31 |

**Table S17. Effects of age, sex, and hindleg-to-weight ratio on travel time to the reward arm during the main escape learning trial.** ANCOVA showed that age group significantly predicted reward arm travel time (R² = 0.325). Using adult females as the reference group, mid-age crickets exhibited longer travel times to the reward arm, while geriatric crickets did not differ. Neither sex nor hindleg-to-weight ratio was associated with travel time. P-values were adjusted using the Holm–Bonferroni method.

| **Source** | **SS** | **df** | **MS** | **F** | ***P*** |
| --- | --- | --- | --- | --- | --- |
| Regression | 742.6 | 4 | 185.7 | 5.01 | **0.0021** |
| Group | 178.6 | 2 | 89.30 | 2.41 | 0.10 |
| Sex | 1.733 | 1 | 1.733 | 0.05 | 0.83 |
| Femoral Volume | 28.64 | 1 | 28.64 | 0.77 | 0.38 |
| Residual | 1595 | 43 | 37.09 |  |  |
| Total | 2337 | 47 |  |  |  |
| **Predictor** | **Estimate** | **Standard error** | | **95% CI** | ***P*** |
| Intercept | 6.98 | 3.17 | | 0.59 to 13.37 | **0.033** |
| Group[Mid-Age] | 6.80 | 3.14 | | 0.47 to 13.13 | **0.036** |
| Group[Geriatric] | 2.02 | 2.26 | | -2.55 to 6.58 | 0.38 |
| Sex[M] | -0.50 | 2.29 | | -5.12 to 4.13 | 0.83 |
| Femoral Volume | 206.9 | 235.4 | | -267.9 to 681.6 | 0.38 |

**Table S18. Effects of age, sex, and femoral volume on travel time to the reward arm during the main escape learning trial.** ANCOVA indicated that the model explained 31.8% of the variance in reward arm travel time (R² = 0.318). Using adult females as the reference group, mid-age crickets took longer to reach the reward arm, while geriatric crickets did not differ. Neither sex nor femoral volume was associated with travel time. *P*-values were adjusted using the Holm–Bonferroni method.

| **Source** | **SS** | **df** | **MS** | **F** | ***P*** |
| --- | --- | --- | --- | --- | --- |
| Regression | 779.9 | 4 | 195.0 | 5.38 | **0.0013** |
| Group | 173.0 | 2 | 86.48 | 2.39 | 0.10 |
| Sex | 0.31 | 1 | 0.31 | 0.01 | 0.93 |
| Femoral CSA | 65.94 | 1 | 65.94 | 1.82 | 0.18 |
| Residual | 1557 | 43 | 36.22 |  |  |
| Total | 2337 | 47 |  |  |  |
| **Predictor** | **Estimate** | **Standard error** | | **95% CI** | ***P*** |
| Intercept | 4.96 | 3.65 | | -2.40 to 12.32 | 0.18 |
| Group[Mid-Age] | 6.25 | 2.86 | | 0.482 to 12.01 | **0.034** |
| Group[Geriatric] | 2.13 | 2.13 | | -2.17 to 6.43 | 0.32 |
| Sex[M] | -0.20 | 2.11 | | -4.45 to 4.06 | 0.93 |
| Femoral CSA | 162.4 | 120.4 | | -80.34 to 405.2 | 0.18 |

**Table S19. Effects of age, sex, and femoral CSA on travel time to the reward arm during the main escape learning trial.** ANCOVA showed that the model accounted for 33.4% of the variance in reward arm travel time (R² = 0.334). Using adult females as the reference group, mid-age crickets required more time to reach the reward arm, while geriatric crickets did not differ. Neither sex nor femoral CSA was associated with travel time. *P*-values were adjusted using the Holm–Bonferroni method.

| **Source** | **SS** | **df** | **MS** | **F** | ***P*** |
| --- | --- | --- | --- | --- | --- |
| Regression | 742.5 | 4 | 185.6 | 5.00 | **0.0021** |
| Group | 202.2 | 2 | 101.1 | 2.73 | 0.077 |
| Sex | 1.59 | 1 | 1.59 | 0.04 | 0.84 |
| Femoral SA/V | 28.46 | 1 | 28.46 | 0.77 | 0.39 |
| Residual | 1595 | 43 | 37.09 |  |  |
| Total | 2337 | 47 |  |  |  |
| **Predictor** | **Estimate** | **Standard error** | | **95% CI** | ***P*** |
| Intercept | 16.51 | 8.35 | | -0.34 to 33.35 | 0.055 |
| Group[Mid-Age] | 6.96 | 3.02 | | 0.88 to 13.04 | **0.026** |
| Group[Geriatric] | 2.08 | 2.24 | | -2.44 to 6.60 | 0.36 |
| Sex[M] | -0.48 | 2.31 | | -5.13 to 4.18 | 0.84 |
| Femoral SA/V | -0.25 | 0.29 | | -0.83 to 0.33 | 0.39 |

**Table S20. Effects of age, sex, and femoral SA/V ratio on travel time to the reward arm during the main escape learning trial.** ANCOVA indicated that the model explained 31.8% of the variance in reward arm travel time (R² = 0.318). Using adult females as the reference group, mid-age crickets took longer to reach the reward arm, whereas geriatric crickets did not differ. Neither sex nor femoral SA/V was associated with travel time. *P*-values were adjusted using the Holm–Bonferroni method.

A.	B.	C.

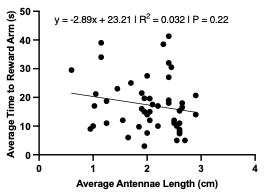

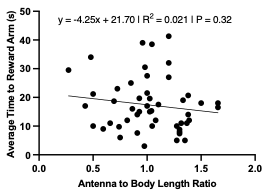

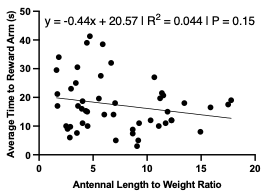

D.	E.	F.

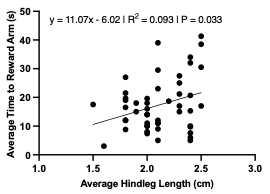

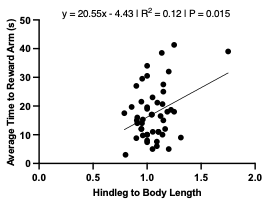

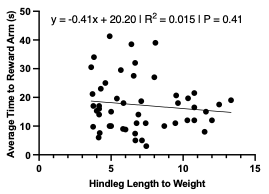

G.	H.	I.

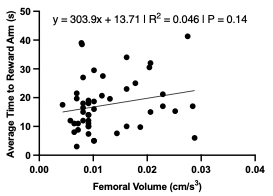

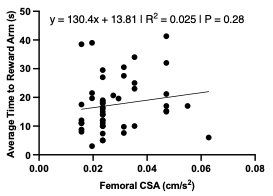

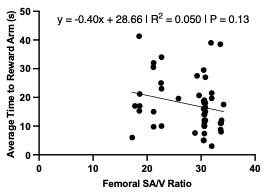

| **Trait** | **β** | **95% CI** | **R²** | ***P*-value** |
| --- | --- | --- | --- | --- |
| Antennal Length (cm) | -2.89 | -7.52 to 1.74 | 0.032 | 0.22 |
| Antennal-to-Body Length | -4.25 | -12.76 to 4.25 | 0.021 | 0.32 |
| Antennal-to-Weight | -0.44 | -1.05 to 0.16 | 0.044 | 0.15 |
| Hindleg Length (cm) | 11.07 | 0.91 to 21.23 | 0.093 | **0.033** |
| Hindleg-to-Body Length | 20.55 | 4.11 to 36.99 | 0.12 | **0.015** |
| Hindleg-to-Weight | -0.41 | -1.39 to 0.57 | 0.015 | 0.41 |
| Femoral Volume (cm^3^) | 303.9 | -108.3 to 716.1 | 0.046 | 0.14 |
| Femoral CSA (cm/s^2^) | 130.4 | -109.9 to 370.8 | 0.025 | 0.28 |
| Femoral SA/V | -0.40 | -0.91 to 0.11 | 0.050 | 0.13 |

**Figure S6. Linear associations between morphological traits and travel time to the reward arm during the test escape learning trial.** Panels **(A–I)** depict simple linear regressions relating reward arm travel time to individual morphological parameters: **(A)** antennal length, **(B)** antennal-to-body length ratio, **(C)** antennal-to-weight ratio, **(D)** hindleg length, **(E)** hindleg-to-body length ratio, **(F)** hindleg-to-weight ratio, **(G)** femoral volume, **(H)** femoral CSA, and **(I)** femoral SA/V ratio. Regression slopes (β), 95% confidence intervals, R² values, and P-values are summarized in the accompanying table. Negative slopes indicate faster travel times with increasing trait size, whereas positive slopes reflect delayed travel times.

| **Source** | **SS** | **df** | **MS** | **F** | ***P*** |
| --- | --- | --- | --- | --- | --- |
| Regression | 1647 | 4 | 411.9 | 7.16 | **0.0002** |
| Group | 1005 | 2 | 502.5 | 8.73 | **0.0006** |
| Sex | 109.7 | 1 | 109.7 | 1.91 | 0.17 |
| Hindleg Length | 29.83 | 1 | 29.83 | 0.52 | 0.48 |
| Residual | 2533 | 44 | 57.56 |  |  |
| Total | 4180 | 48 |  |  |  |
| **Predictor** | **Estimate** | **Standard error** | | **95% CI** | ***P*** |
| Intercept | 2.818 | 14.15 | | -25.69 to 31.33 | 0.84 |
| Group[Mid-Age] | 10.56 | 3.67 | | 3.16 to 17.96 | **0.0062** |
| Group[Geriatric] | -1.93 | 2.93 | | -7.84 to 3.98 | 0.51 |
| Sex[M] | 3.71 | 2.69 | | -1.71 to 9.12 | 0.17 |
| Hindleg Length | 4.92 | 6.84 | | -8.86 to 18.71 | 0.48 |

**Table S21. Effects of age, sex, and hindleg length on travel time to the reward arm during the test escape learning trial.** ANCOVA demonstrated that age group significantly predicted reward arm travel time (R² = 0.394). Using adult females as the reference group, mid-age crickets required more time to reach the reward arm, whereas geriatric crickets did not differ. Neither sex nor hindleg length was associated with variation in travel time. *P*-values were adjusted using the Holm–Bonferroni method.

| **Source** | **SS** | **df** | **MS** | **F** | ***P*** |
| --- | --- | --- | --- | --- | --- |
| Regression | 1741 | 4 | 435.2 | 7.85 | **<0.0001** |
| Group | 1226 | 2 | 613.1 | 11.06 | **0.0001** |
| Sex | 79.95 | 1 | 79.95 | 1.44 | 0.24 |
| Hindleg-to-Body Length | 123.2 | 1 | 123.2 | 2.22 | 0.14 |
| Residual | 2439 | 44 | 55.44 |  |  |
| Total | 4180 | 48 |  |  |  |
| **Predictor** | **Estimate** | **Standard error** | | **95% CI** | ***P*** |
| Intercept | 1.92 | 7.64 | | -13.48 to 17.32 | 0.80 |
| Group[Mid-Age] | 11.09 | 2.78 | | 5.49 to 16.69 | **0.0002** |
| Group[Geriatric] | -1.21 | 2.54 | | -6.32 to 3.90 | 0.64 |
| Sex[M] | 2.59 | 2.15 | | -1.75 to 6.93 | 0.24 |
| Hindleg-to-Body Length | 10.79 | 7.24 | | -3.80 to 25.38 | 0.14 |

**Table S22. Effects of age, sex, and hindleg-to-body length ratio on travel time to the reward arm during the test escape learning trial.** ANCOVA showed that age group significantly predicted reward arm travel time (R² = 0.417). Using adult females as the reference group, mid-age crickets took longer to reach the reward arm, while geriatric crickets did not differ. Neither sex nor hindleg-to-body length ratio was associated with travel time. *P*-values were adjusted using the Holm–Bonferroni method.
