## Supplementary material for "Age-related cognitive decline in house crickets reveals conserved patterns of sensory and learning deficits across the lifespan": BioRender Publication Licenses: 3 - Publication License Jul-30-2025.pdf

49 Spadina Ave. Suite 200
Toronto ON M5V 2J1 Canada
www.biorender.com

### Confirmation of Publication and Licensing Rights

**July 30th, 2025**

***Subscription Type:*** *Student Plan - Academic*
***Agreement number:*** *PM28KMPKI6*
***Publisher Name:*** *GeroScience*

***Citation to Use:*** *Created in BioRender. Liao, G. (2025) https://BioRender.com/rb2sq6u*

To whom this may concern,

This document is to confirm that Gerald Liao has been granted a license to use the BioRender Content, including icons, templates, and other original artwork, appearing in the attached Completed Graphic pursuant to BioRender's Academic License Terms. This license permits BioRender Content to be sublicensed for use in publications (journals, textbooks, websites, etc.).

All rights and ownership of BioRender Content are reserved by BioRender. All Completed Graphics must be accompanied by the following citation: "Created in BioRender. Liao, G. (2025) https://BioRender.com/rb2sq6u".

BioRender Content included in the Completed Graphic is not licensed for any commercial uses beyond use in a publication. For any commercial use of this figure, users may, if allowed, recreate it in BioRender under an Industry BioRender Plan.

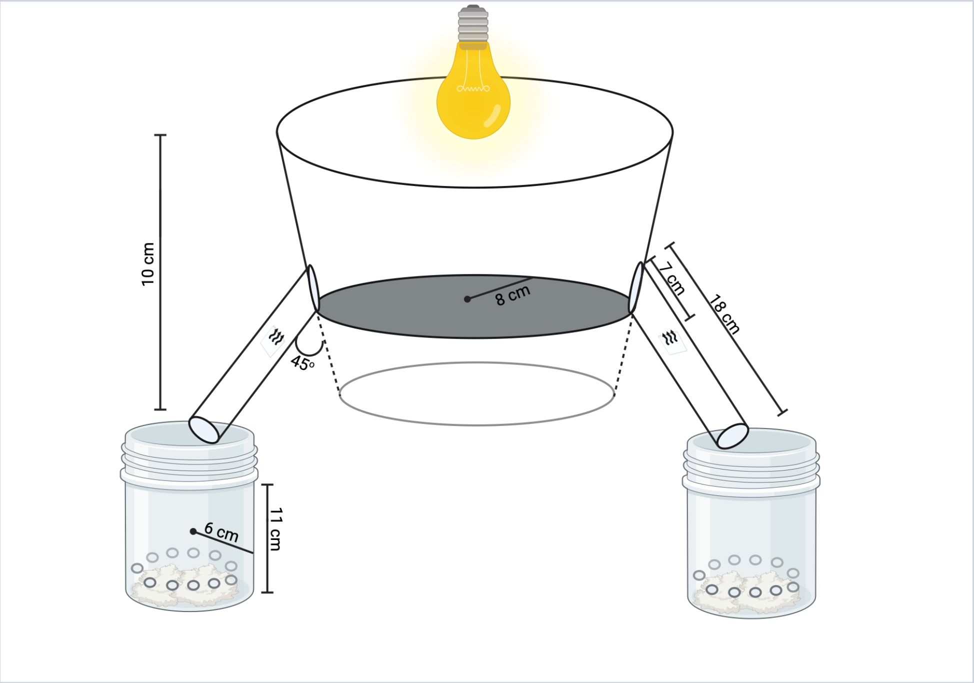


*For any questions regarding this document, or other questions about publishing with BioRender, please refer to our BioRender Publication Guide, or contact BioRender Support at.*
